# Vegetated urban spaces increase *Aedes albopictus* survival and the risk of Dengue and Chikungunya Transmission: a field and modelling study in Montpellier, France

**DOI:** 10.64898/2026.03.02.709080

**Authors:** Colombine Bartholomée, Clémence Garcia-Marin, Céline Sutter, Florence Fournet, Emilie Bouhsira, Nicolas Moiroux

## Abstract

**Introduction:** By 2025, *Aedes albopictus* had spread across France, leading to a rise in indigenous arboviral cases since 2021. While urban greening is promoted for its health and climate benefits, its potential adverse effects on arboviral risk remain understudied. Urban green spaces may enhance mosquito habitats and human-vector contact.

**Methods:** This study investigates the variation in *Aedes albopictus* longevity across months and urban environments in Montpellier, a green Mediterranean city, and its impact on the theoretical basic reproductive number (R_0_) for Chikungunya, Dengue, and Zika viruses. From May to October 2023, female mosquitoes were sampled monthly using CO_2_ traps in three urban environments: urban parks, impervious areas, and residential areas composed of houses with gardens. Daily mosquito survival was estimated from parity rate measurements, and the associated environmental determinants were analysed. Modelled predictions, crude field estimates of vector density and parity rates, and microclimatic data were used to parameterize and feed R_0_ models. For each environment (or sampling area), 1000 monthly R_0_ simulations were run, followed by sensitivity analyses.

**Results/Discussion:** The highest mean daily survival (0.917) was observed in residential areas, compared to parks (0.887) and impervious areas (0.873). A daily average exposure of 10% of possible bites induces potential transmission risks for CHIKV and DENV. R_0_ varied by environment and month, with higher values in residential areas and with greater variability in impervious areas. Parity rate emerged as the main driver of R_0_ variability. These results highlight the value of fine-scale field data for arboviral risk assessment.

## Introduction

Since 2007, Europe has experienced a growing number of local transmissions of *Aedes*-borne arboviruses, including dengue virus (DENV), chikungunya virus (CHIKV), and Zika virus (ZIKV) [1]. These events are linked to the establishment and spread of *Aedes albopictus*, commonly known as the tiger mosquito. Metropolitan France is particularly at risk, reporting the highest number of imported dengue and chikungunya cases in Europe [1], as well as the highest number of local DENV transmission events. More than 80% of all local arboviral cases reported in France since 2006 were recorded after 2021. This may be partly explained by recurrent arboviral outbreaks in the French overseas territories, such as the 2025 dengue epidemic in Guadeloupe [2].

Since 2020, the European Commission, as part of the Green Deal, has outlined 100 actions to be implemented by 2030 to mitigate the effects of climate change and urbanisation. One of these actions is the promotion of urban greening in cities with population over 20 000 [3]. The Occitanie region has implemented multiple urban greening initiatives. Despite its many benefits, concerns arose in 2024 about potential adverse effects related to vectors and vector-borne diseases [4]. A recent scoping review revealed that the impact of urban greening on vectors varies depending on the vector system studied and the local context, with most studies focusing on *Aedes albopictus* or dengue transmission risk and reporting harmful correlations with urban vegetation indicators [5]. We recently conducted the first study on this topic in France, showing that urban vegetation promotes the abundance of *Ae. albopictus* [6].

However, an increase in abundance alone is insufficient to assess the influence of urban vegetation on transmission risk [7]. In particular, vector survival is a key determinant of transmission, as only mosquitoes that live long enough can complete the extrinsic incubation period and become infectious. Urban vegetation may strongly affect this parameter by providing cooler, more humid shelters and increasing the availability of blood and sugar sources, thereby enhancing mosquito survival compared to more artificial environments. Conversely, vegetation may also promote greater biodiversity, potentially increasing the abundance of mosquito predators, which could reduce mosquito survival.

In this study, we proposed to study the environmental determinants of *Ae. albopictus* parity rates (a proxy of survival) in the city of Montpellier, the greenest in the Occitanie region, France. We then integrated field-derived estimates of survival and abundance into an *R*_*0*_ model to assess the risk of local transmission for DENV, CHIKV, and ZIKV over six months and across three urban environments with varying vegetation coverage.

## Methods

### Study design and entomological data collection

This study was conducted from May to October 2023 in Montpellier. We sampled nine areas across three urban environments with varying degrees of vegetation cover. These areas were two urban parks (PRK-1 and PRK-2), three residential neighbourhoods composed of houses with private gardens (RES-1, RES-2, and RES-3), and four densely built impervious environments (IMP-1, IMP-2, IMP-3, and IMP-4). Each mosquito sampling session lasted 48 hours and was conducted once per month. Mosquitoes were retrieved from each BG-PRO trap every 24 hours. The study design and entomological data collection were extensively described in a previous study by the same authors [6]. In order to further analyse mosquito survival, we used the Detinova method [8] to dissect the ovaries of females *Ae. albopictus* and determine whether they were parous or nulliparous, based on the appearance of the ovarian tracheoles (Supplementary Figure S1). Gravid females (with eggs) were excluded from the subsequent analysis, as it is impossible to ascertain whether they had already completed a gonotrophic cycle.

### Environmental, demographic, and microclimatic data collection and preparation

A fine-scale land cover map of Montpellier was created. Various landscape indices and human population densities were calculated for multiple buffer sizes around each trap. Fine-scale population data was obtained at the level of living areas, and population counts were summed for different buffer sizes. Hourly temperature and relative humidity data were collected continuously using Hygro Button data loggers placed near each trap. For each sampling session, daily minimum, maximum, and average values for both temperature and relative humidity were calculated over 24-hours periods. The collection, processing, and preparation of environmental and microclimatic data have been previously described [6].

### Data analysis

#### *Environmental determinants of* Ae. *albopictus survival*

To assess environmental determinants of survival, we analysed parity rates of *Ae. albopictus* females collected in the traps. Parity rate is defined as the proportion of parous female mosquitoes in a population, i.e. those that have completed at least one gonotrophic cycle and laid eggs. Parity rate is closely related to the daily survival rate (Supplementary Table S1). Parity rates were calculated for each trap over 24-hour sampling periods. Bivariate analyses were performed to assess associations between parity rates and environmental, microclimatic, and demographic variables. Parity rates per trap per 24h were modelled using generalised linear mixed models (GLMMs) with binomial error distribution and random intercepts for sampling sessions, weighted by the corresponding number of dissected females (parous and nulliparous). To compare parity rates specifically between sampling environments, an additional GLMM was fitted, including only the sampling environment as a fixed effect. The selection of explanatory variables and exclusion of collinear predictors using the variance inflation factor followed a previously described procedure [6]. A multivariable GLMM was subsequently performed using the retained predictors, with stepwise selection based on the Akaike Information Criterion (AIC). Temporal and spatial autocorrelation of the residuals was assessed. All analyses were conducted in R version 4.2.3, using the packages ‘glmmTMB’ [9], and ‘performance’ [10].

#### Predictive models of parity rate and female abundance per trap per day

A predictive binomial generalised additive model (GAM) was fitted to parity rates measured in each trap per 24-h period, weighted by the corresponding number of dissected females. Explanatory variables included the sampling area and the month of collection, the latter modelled using a thin plate regression spline, with a random intercept for the sampling session. This smoother allows flexible modelling of non-linear relationships without requiring manual knot placement [11]. GAMs were chosen for these predictive models because they can capture complex seasonal and environmental patterns without requiring a priori specification of functional forms, making them particularly suitable for generating realistic predictions to parameterize the *R*_*0*_ model (see below). A second predictive GAM was fitted to the number of females caught per trap per day (the “abundance” model), using a negative binomial distribution and including a random intercept for the sampling session. The same explanatory variables were used as in the parity rate GAM. Both GAMs were fitted in R using the ‘mgcv’ package [12].

#### Calculation and simulation of the basic reproduction number (R_0_) using field or GAM-predicted data

Local transmission risk was assessed using *R*_*0*_, defined as the expected number of secondary human cases generated by introducing a single infected individual into a fully susceptible population. When *R*_*0*_>1, local transmission is possible, and the disease may spread in the absence of control; when *R*_*0*_<1, transmission is unlikely [13]. We used a classical formula of *R*_*0*_ derived from the Ross-MacDonald model [7]:

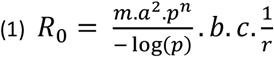

where *m. a* is the human biting rate, *a* the human feeding rate, *p* the daily survival rate of mosquitoes, *n* the duration of the extrinsic incubation period of the virus, *b* infectivity of mosquito to human, *c* infectivity of human to mosquito, and 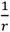 the duration of the human infectious period.

Detailed definitions of parameters, their values or formulae, and associated references are provided in Supplementary Table S1.

We did not use the total number of *Ae. albopictus* females caught or predicted per trap per day as a direct proxy for the human biting rate *m.a*. Because people are not continuously exposed to mosquitoes throughout the day, this approach would have implied unrealistically high levels of human exposure. Methods exist to estimate biting exposure more accurately, but they require detailed data on both human and vector behavior [14,15]. In the absence of such data, we considered three exposure scenarios corresponding to 10%, 20%, and 30% of the mosquito abundance measured or predicted in traps (i.e. *m.a* = *N.Π*, with N the daily abundance per trap and *Π* the exposure level). At 10% exposure (Π=0.1), the average number of bites per human per day was estimated at 0.45 in impervious areas, 0.48 in residential areas, and 0.81 in urban parks, doubling and tripling under the 20% and 30% exposure scenarios, respectively, and considering average crude field estimates of the abundance *N*. All calculations and simulations of *R*_*0*_ were based on these three scenarios. Corresponding monthly variations in daily human exposure to bites by the environment and sampling area are presented in Supplementary Figures S2 and S3. The daily survival rate (*p*) was estimated as the *g*^th^ root of the parity rate (*Par*), i.e. *p=Par*^*1/g*^, where *g* is the duration (in days) of the gonotrophic cycle [16] (Supplementary Table S1). Values of *Par* and *N* parameters were derived from either crude field estimates or the GAM-derived predictions. Field data provide direct empirical measurements, while GAM predictions smooth over sampling variability and capture seasonal and environmental patterns, thus offering continuous estimates. Both approaches allowed a robust assessment of transmission risk. Parameters *g, n*, and *b* (for DENV and ZIKV) were modelled as functions of field-recorded temperature (see microclimatic data above), using formulae from the literature, as detailed in Supplementary Table S1.

*R*_*0*_ was estimated for DENV, CHIKV, and ZIKV, for each month and sampling environment (urban park, impervious areas, and residential areas with private gardens), as well as for each individual sampling area. Uncertainty in the GAM parameter estimates was propagated into the *R*_*0*_ simulations through parametric resampling (1000 simulations per sampling area and month), allowing us to derive empirical 95% confidence intervals for transmission risk.

All scripts and data are available on [10.5281/zenodo.15854094] and on GBIF [17], respectively. An online R shiny application has been developed to better visualise the different results [https://cbartholomee.shinyapps.io/r0_modelling_montpellier/].

#### Sensitivity analysis

A sensitivity analysis of the *R*_*0*_ model was performed using Sobol indices to assess the contribution of each parameter, individually or in interaction, to the overall variability of the outcome, using Monte Carlo methods [18]. This sensitivity analysis was conducted only for DENV, using the formulation for *n* and *b* employed for *R*_*0*_ estimates, with realistic parameter ranges for each parameter. The Sobol indices were computed using the ‘sensitivity’ package [19]. We also assessed the robustness of our results to the choice of *R*_*0*_ formulation by applying an alternative equation used by Poletti *et al*. (2011) [20].

## Results

### Explanatory model of the parity rate

A total of 1106 females were collected (Table 1), of which 1074 were dissected, and 32 were too damaged for dissection. Among those dissected, 125 were gravid and 124 had undetermined parity status (NPI). Parity rate in residential areas was 65.5% (Table 1), significantly higher than 54.0% in impervious areas (Odds-Ratio (OR) [95% Confidence interval (CI)] = 1.61 [1.06; 2.44], p = 0.023), and 52.9% in urban parks (OR [95% CI] = 1.63 [1.17; 2.25], p = 0.003). Parity rates were similar between impervious areas and urban parks (OR [95% CI] = 0.99 [0.68; 1.44], p = 0.96). Assuming that gonotrophic cycle duration is temperature-dependant (Supplementary Table S1), the estimated average daily survival rate was 0.92 in residential areas compared to 0.887 in urban parks and 0.873 in impervious areas (Table 1).

**Table 1.** Summary of entomological data, local temperature and derived survival rates by sampled environment, aggregated across all months. The average local temperature was calculated using field microclimatic data. * based on temperature, according to a regression model fitted on [21] data, see formulae in Supplementary Table S1, **\*\*** based on *g* and parity rate, see formulae in Supplementary Table S1. ND = Not determinated

| Environment | N females | N dissected | N ND | N gravids | N nulliparous (a) | N parous (b) | Parity rate = $b/(a+b)$ (Par) | Average local Temperature | Average duration of gonotrophic cycle (in days)* (g) | Average daily survival rate** $p = Par^{1/g}$ |
| --- | --- | --- | --- | --- | --- | --- | --- | --- | --- | --- |
| Impervious | 234 | 221 | 30<br>(30/221,<br>13.6%) | 41 | 69 | 81 | 0.540 | 24.25°C | 4.53 | 0.873 |
| Urban parks | 525 | 519 | 52<br>(52/519,<br>10.0%) | 47 | 198 | 222 | 0.529 | 23.03°C | 5.33 | 0.887 |
| Residential areas | 347 | 334 | 42<br>(32/334,<br>12.6%) | 37 | 88 | 167 | 0.655 | 23.64°C | 4.91 | 0.917 |

After bivariate analysis (Supplementary Figures S4, S5, and S6), multicollinearity assessments, and stepwise selection based on AIC, four variables were retained for the multivariate GLMM of parity (reasons for exclusion are provided in Supplementary Table S2). Three variables were significantly and positively associated with parity: the average size of low vegetation patches (in m^2^) within 100 m of the trap (OR [95% CI] = 1.003 [1.001; 1.005], p = 0.009), the total building perimeter (in m) within 20 m (OR = 1.004 [1.001; 1.007], p = 0.008), and the maximum relative humidity (in %) recorded during sampling (OR = 1.021 [1.005; 1.037], p = 0.008).

### Analysis of *R*_*0*_ based on field observations and GAM predictions

*R*_*0*_ simulations indicated that the risk of CHIKV and DENV transmission is not negligible (upper 95% confidence limit >1) even at 10 % of bite exposure, across all environments from June to October for CHIKV, and in urban parks and residential areas during June/July and September for DENV (Figure 1). The risk of ZIKV transmission is negligible across all tested human exposure scenarios. *R*_*0*_ values varied over time and environments: estimations based on crude field estimates of parity rate and abundance showed bimodal patterns, with peaks in June–July and September–October depending on the environment. The same temporal trend was observed with simulations based on GAM predictions. *R*_*0*_ values derived from GAM predictions were generally lower than those derived from field estimates (Figure 2). Across all viruses and bite exposure levels, residential areas showed the highest *R*_*0*_ values (Figures 1 and 2). For example, for DENV, under the 20% exposure scenario, the average *R*_*0*_ derived from field estimates was 1.57 in impervious areas, 1.77 in urban parks, and 2.31 in residential areas (Supplementary Table S3). The corresponding average *R*_*0*_ values based on GAM predictions were 0.18 (95 % CI = [0.07; 0.74]), 0.29 (95 % CI = [0.11; 1.27]), and 0.52 (95 % CI = [0.19; 2.16]), respectively. Additionally, average *R*_*0*_ values based on GAM predictions remained above 1 for one month in residential areas, whereas in urban parks and impervious areas, they never exceeded 1 (Figure 1, Supplementary Table S4).

**Figure 1.**
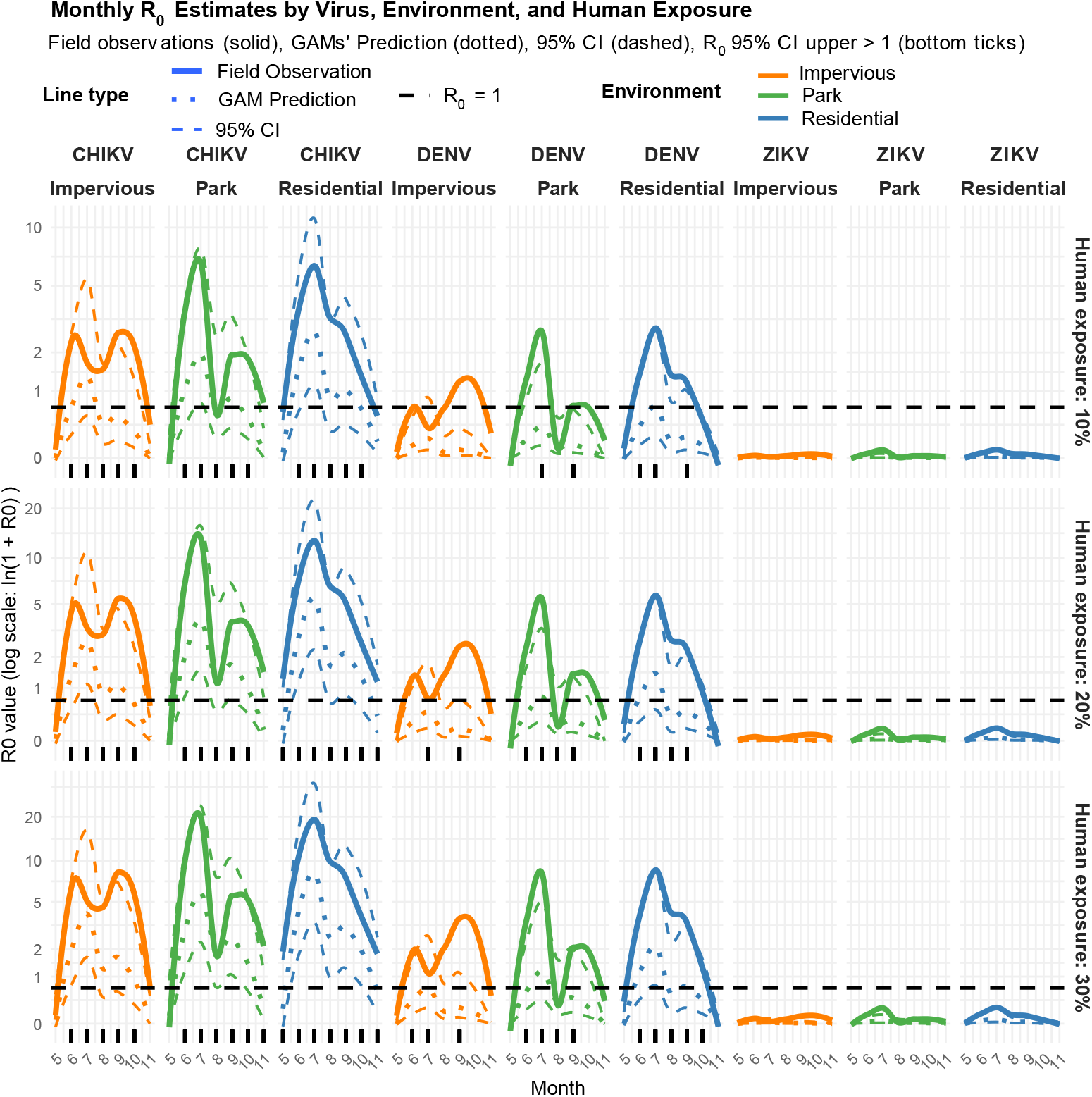
Monthly *R*_*0*_ estimates based on crude field averages or GAM-predicted parity rates and abundance, across different environments, viruses (CHIKV, DENV, and ZIKV), and three levels of human exposure in Montpellier, France, 2023. *R*_*0*_ according to the Ross-MacDonald formulae. The solid line represents *R*_*0*_ derived from crude field estimates of parity rates and abundance, the dotted line corresponds to *R*_*0*_ simulated from GAM predictions, and the dashed lines indicate the 95% confidence intervals of *R*_*0*_ simulations. The horizontal black line denotes the epidemic threshold (*R*_*0*_ = 1). Thickened ticks along the x-axis indicate months when the upper 95% confidence limit of *R*_*0*_ simulations exceeded 1. The y-axis is on log-scale.

**Figure 2.**
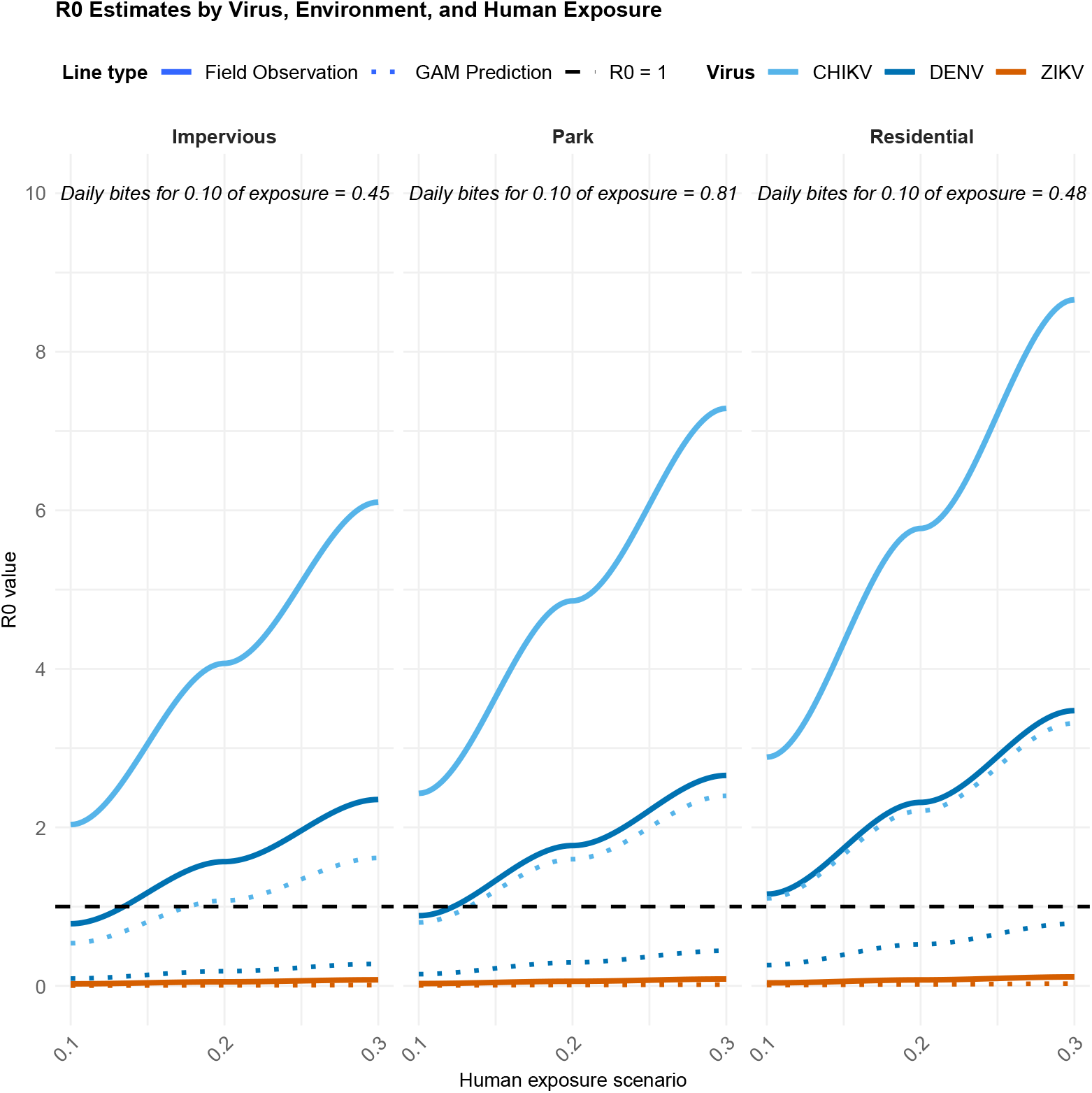
Average *R*_*0*_ for three arboviruses according to the environment and the level of human exposure to bites in Montpellier (France), in 2023. The solid lines show *R*_*0*_ estimates derived from crude field estimates of parity rates and abundance. The dotted lines show *R*_*0*_ simulated from GAM predictions. The horizontal black line indicates the epidemic threshold (*R*_*0*_ = 1). Human exposure scenarios represent the proportion of the average daily mosquito catch assumed to bite a single individual.

Monthly *R*_*0*_ estimates revealed distinct temporal patterns across sampling areas (Figure 3). The *R*_*0*_ values derived from field estimates and those from GAM predictions generally followed similar seasonal trends, although some differences were observed in IMP-1, IMP-2, IMP-3, and RES-2. Greater variability in *R*_*0*_ values was observed in impervious areas than in residential areas and in urban parks. For instance, for CHIKV, at a human exposure level of 10%, the upper confidence interval of *R*_*0*_ simulations from GAM prediction exceeded 1 from June to October in IMP-1, with a maximum upper bound of 10.49. In contrast, at IMP-3, *R*_*0*_ simulations from GAM prediction remained below 1, with a maximum upper bound of 0.421—approximately twenty-five times lower than in IMP-1.

**Figure 3.**
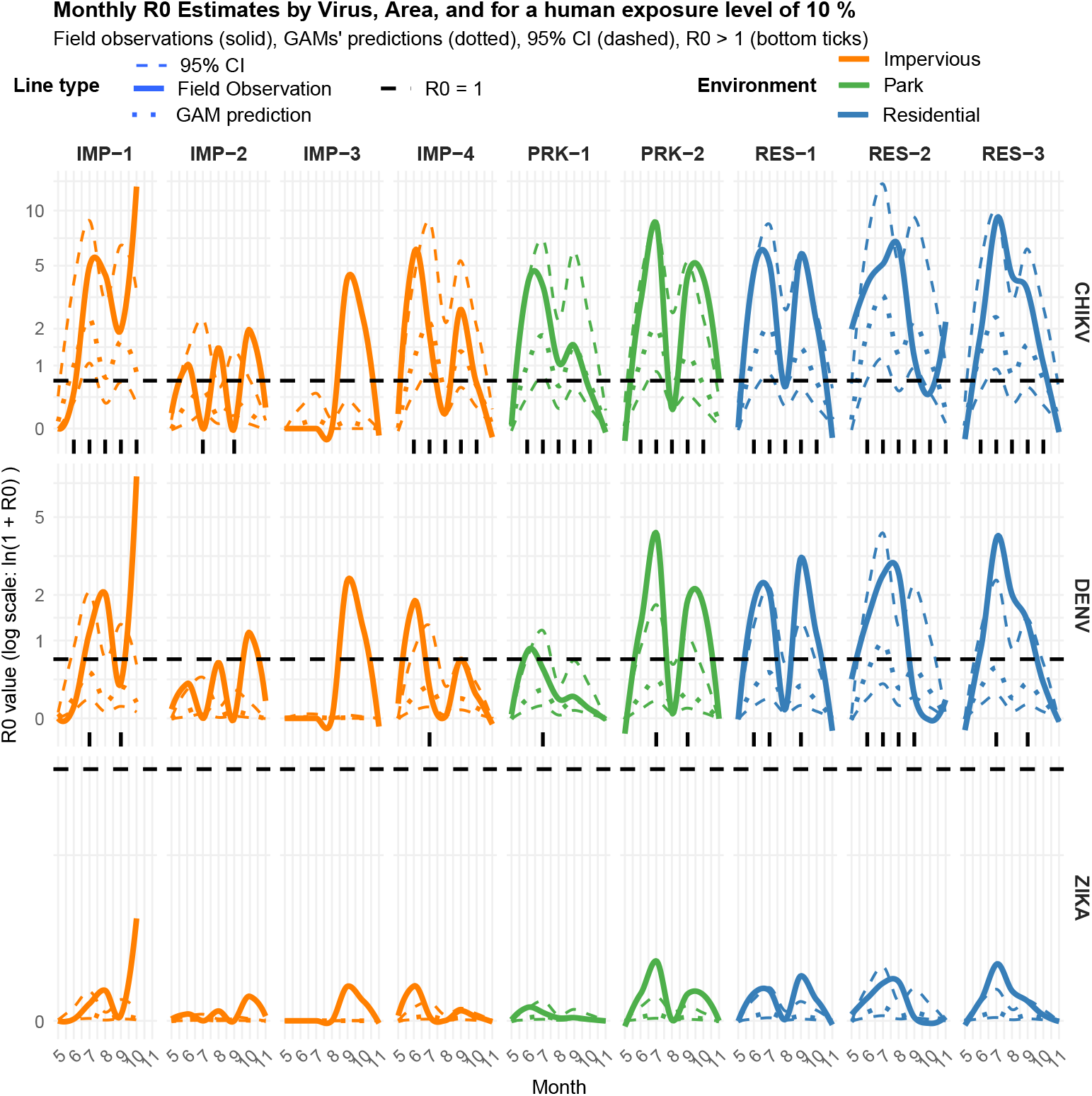
Monthly *R*_*0*_ estimates derived from crude field averages and GAM-predicted parity rates and abundance, across sampling areas and viruses (CHIKV, DENV, and ZIKV), assuming a 10% human exposure level, Montpellier, France, 2023. The solid line shows the *R*_*0*_ derived from crude field estimates of parity rates and abundance, the dotted line corresponds to *R*_*0*_ simulated from GAM predictions, and the dashed lines indicate the 95% confidence intervals of *R*_*0*_ simulations. The horizontal black line denotes the epidemic threshold (*R*_*0*_ = 1). Thickened ticks along the x-axis indicate months when the upper 95% confidence limit of *R*_*0*_ simulations exceeded 1. The y-axis is on log-scale.

### Sensitivity analysis of *R*_*0*_ model

The sensitivity analysis using Sobol indices revealed that *R*_*0*_ outcomes were highly sensitive to the parity rate (Figure 4), with a first-order Sobol index exceeding 0.35 and a total-order index (taking interactions into account) above 0.75. Model outcomes were also significantly sensitive to three other parameters: the number of females caught per trap per day (*N*), the human exposure to bite (*Π*), and the infectivity of human to mosquito (*c*).

**Figure 4.**
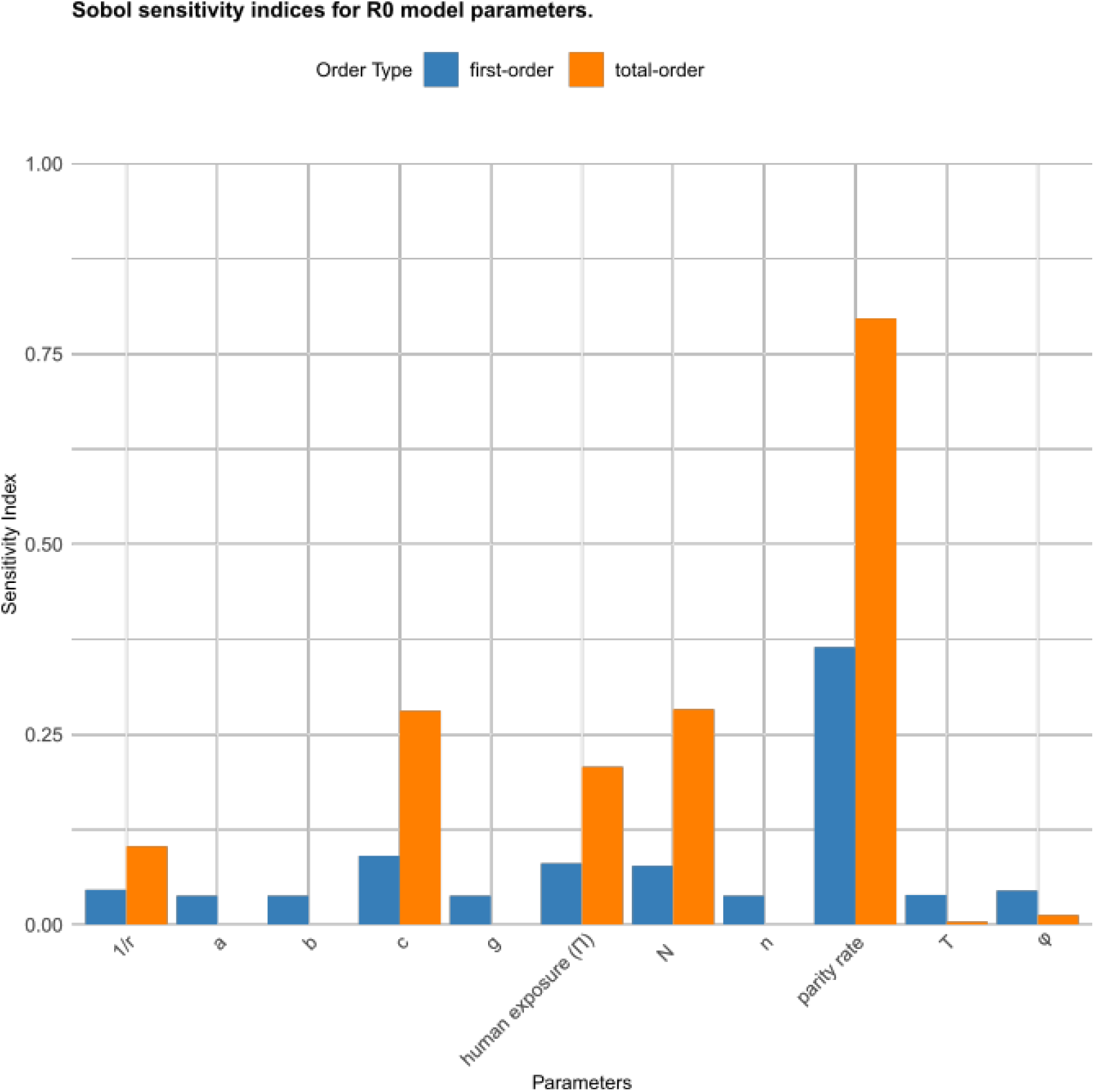
Sobol sensitivity indices for DENV *R*_*0*_ model parameters. Parameters: *1/r* = duration of the human infectious period, *a* = number of blood meals taken from humans per mosquito per day, *b* = vector competence calculated using the formula from Benkimoun *et al*. (2021) [22], *c* = host competence, *g* = gonotrophic cycle duration, *Π =* human exposure, *N* = number of females caught per trap per day, *n* = extrinsic incubation period calculated using the formula from Caminade *et al*. (2017) [23], *T* = local temperature and φ = human preference index. First-order and total-order Sobol indices indicate the main and interaction effects of each parameter on the variance of R_0_ estimates.

## Discussion

In this study, we successfully calculated monthly *R*_*0*_ values from field-collected entomological and microclimatic data at a fine spatial scale. Both the parity rate and *R*_*0*_ estimates were higher in residential areas than in impervious areas and urban parks. *R*_*0*_ exhibited marked spatiotemporal variability, mostly driven by parity rates and abundance of females, level of human exposure, and infectivity of human to mosquito.

Average parity rates in field-collected *Ae. albopictus* females ranged from 52.9% to 65.5%, with the highest values in residential areas. This pattern is consistent with findings by Delatte *et al*. (2013) in La Réunion Island, where parity rates ranged from 63.0% to 76.5% and were similarly higher in residential areas [24]. Moreover, we found that parity rates were positively influenced by low vegetation within a 100 m buffer and by building length within a 20 m radius around traps, both of which are characteristic of residential areas. We hypothesized that low vegetation provided suitable resting sites, while greater building length may reflect higher availability of blood meals. Consequently, residential areas exhibited higher arbovirus transmission risk, as indicated by the *R*_*0*_ model, which may explain why past local transmission events were predominantly recorded in such environments [25]. For example, of the 34 locally acquired chikungunya cases reported in Hérault (the department of Montpellier, in Occitanie), 29 occurred in four towns near Montpellier that are mainly residential areas with private gardens: Prades-le-Lez (1 case), Castries (16 cases), Mauguio (11 cases), and Candillargues (1 case).

Simulated CHIKV *R*_*0*_ values with a human exposure level of 10% highlighted a risk of local transmission during almost all months of mosquito activity and all environments, while for DENV the risk of local transmission was restrained to July and September in urban parks and residential areas. Risk for local transmission of ZIKV was always negligible. These differences can be explained by the shorter extrinsic incubation period (n) of CHIKV compared with DENV and ZIKV (Supplementary Figure S13) and lower vector competence of *Ae. albopictus* for ZIKV than for CHIKV and DENV (Supplementary Figure S14) [26]. *R*_*0*_ estimates followed a bimodal distribution, with peaks occurring in June–July and in September– October. Comparable seasonal patterns were reported by Solimini *et al*. (2018) in Roma (Italy), who also observed higher transmission risks for CHIKV and DENV compared to ZIKV, with two seasonal peaks in August and October [27].

However, *R*_*0*_ values derived from our field data were higher than those reported by Solimini *et al*. (2018) [27]. As noted by Liu *et al*. (2020), *R*_*0*_ can vary substantially depending on climate and methodology [28]. In temperate regions, average *R*_*0*_ values were 1.88 for CHIKV, 3.69 for DENV, and 0.53 for ZIKV [28], compared with 0-13.32, 0-7.62, and 0.02-0.04, respectively, in our study with 10% of human exposure to bite. These differences in *R*_*0*_ values reflect methodological choices. Solimini *et al*. (2018) estimated the human biting rate by multiplying the mosquito biting rate by the vector-to-host ratio, which was obtained by dividing the estimated uniform vector density by the assumed uniform human density [27]. In our study, we observed differences in mosquito abundance across urban environments but we did not take human density into account and we assumed that the parameters *m. a* could be approximated by the number of females/trap/day, based on the hypothesis that one BG-Pro trap is equivalent to one immobile human host. This assumption is supported by a study in China, which reported no significant differences between female mosquito abundance in BG-Sentinel traps and human landing catches [29]. However, no equivalent validation study exists in France.

Furthermore, female abundance was defined as the number of mosquitoes caught per trap over a 24-hour period, which may not accurately reflect realistic human exposure scenarios. This is the reason why we used different scenario of human exposure. To improve model accuracy, a behavioural survey investigating human presence in various urban environments, exposure times, and the use of personal protection measures against mosquito bites should be conducted. Other useful data would include the hourly biting activity of *Ae. albopictus* in different urban environments and months. Finally, in our study, between 10% and 13.6% of dissected females had an undetermined parity status, which may affect the estimated parity rates and *R*_*0*_. This observation is consistent with Joy *et al*. (2012), where approximately 10% of *Ae. aegypti* females also had undetermined status [30].

It is also important to note that the *R*_*0*_ estimates were derived from entomological data collected in specific urban areas, with sampling conducted twice a month for six months in 2023. These *R*_*0*_ estimates are inherently connected to the specific environmental and temporal conditions. The *R*_*0*_ values we obtained should be interpreted more as infection potential indicators rather than absolute reproductive numbers. They serve as proxies and should not be extrapolated beyond the context of the study. They should be interpreted as part of an integrated framework to identify spatial and temporal hotspots of transmission risk.

Our study introduces valuable indicators (parity rates and abundance) to assess the relative risk of local transmission. The methodology developed allows for the incorporation of local microclimatic temperatures and urban environmental characteristics in the estimation of *R*_*0*_. This enables accounting for the differential influence of temperature on key entomological and viral parameters, and the identification of specific periods and urban areas where transmission risk is elevated. The sensitivity analysis revealed that the main contributors to *R*_*0*_ variability was vector longevity. This is consistent with the structure of the *R*_*0*_ formula [7], in which the parity rate has an exponential effect. This indicates that vector control strategies targeting vector longevity should be prioritize during epidemics events.

## Collaborators

Non applicable.

Statements

### Ethical statement

This study involves the collection of mosquitoes (Aedes albopictus) in urban settings, which does not require specific approval under the NAGOYA Protocol or ethical approval.

### Use of artificial intelligence tools

ChatGPT was only used for minor rephrasing.

### Data availability

Entomological data are available on the GBIF [https://www.gbif.org/dataset/8e52f35a-2522-4865-8361-5c249310a7cf].

R scripts are available on [10.5281/zenodo.15854094].

An online R shiny application has been developed to better visualise the different results [https://cbartholomee.shinyapps.io/r0_modelling_montpellier/].

### Authors contribution

CB, NM and FF were involved in designing the sampling plan. CB collected the field data. CB, CS and CGM dissected female mosquitoes. CGM and CB performed the multivariate analysis of the parity rate. CB and NM modelled the R0 and developed the R scripts. CB wrote the manuscript, which was then corrected by NM, EB and FF. Every co-author revised the manuscript.

## Acknowledgements

We would like to thank Gilbert Le Goff, Nil Rahola and Christophe Paupy for their help with the sampling design, and Mathilde Mercat and Coralie Grail for their help in the field and with data collection. Our thanks also go to the managers and staff of the different sampling sites. Finally, we would like to thank the Défi Clé RIVOC initiative, the Occitanie Region, and the University of Montpellier for their funding.

## Abbreviation

AIC: Akaike Information Criterion
CHIKV: Chikungunya Virus
DENV: Dengue Virus
GAM: Generalized Additive Model
GLMM: Generalized Linear Mixed Models
ZIKV: Zika Virus

