## Supplementary files for "Vegetated urban spaces increase *Aedes albopictus* survival and the risk of Dengue and Chikungunya Transmission: a field and modelling study in Montpellier, France"

### **Additionnal files**

Content

#### Supplementary text

**Supplementary Text S1: Methodology used to estimate the basic reproduction number R_0_**

To estimate the R_0_, two formula were used:

(1) the first, proposed by Ross-MacDonald [1]:

$$R_{0}= \frac{m\times a^{2}\times p^{n}}{-\log\left( p \right)} \times b\times c \times\frac{1}{r}$$

(2) The second, from Poletti *et al* [2]:

$$R_{0}=R_{0}^{HV} \times R_{0}^{VH}=\left( \frac{M}{H} \times a \times c \times\frac{1}{r} \times\frac{\frac{1}{n}}{\frac{1}{n}+\tau} \right) \times\left( a \times b x \frac{1}{\tau} \right)$$

Where:

- $R_{0}^{HV}$ is the number of vectors directly infected by the introduction of a single infectious host into a fully susceptible vector population.
- $R_{0}^{VH}$ is the number of hosts directly infected by the introduction of a single infectious vector into a fully susceptible host population.
- $\frac{M}{H}$ is the ratio of vector (female) density to host (human) density, approximated in our study by $m$.
- $\tau$ is the vector mortality rate, approximated as $-\log(p)$, where $p$ is the daily survival rate.

The definitions, values, and references for all parameters used in the models are provided in the additional table 1.

#### Supplementary tables

**Supplementary Table 1: Parameter equations and values used to estimate R₀ for DENV, CHIKV, and ZIKV**

| **Parameters and variables** | **Description** | **Virus** | **Source of data, value or Equation** | **Range of value used in sensitivity analysis** | **Reference** |
| --- | --- | --- | --- | --- | --- |
| $m.a$ | Human biting rate: number of *Ae. albopictus* bites per human per day | - | $ma=N.\Pi$ |  | - |
| $N$ | Number of *Ae. albopictus* females per trap per 24h | - | Crude field data or prediction of the ‘abundance’ GAM model | [0; 65] | - |
| $\Pi$ | Exposure index: proportion of trapped female a human is expected to be exposed to | - | 0.1, 0.2 or 0.3 | [0.1; 1] | - |
| $a$ | Human feeding rate: number of blood meals taken from humans per mosquito per day | - | $a=\frac{\varphi}{g}$ | [0.078; 0.30] | [3] |
| $\varphi$ | Human preference index: proportion of blood meal taken on humans | - | 0.842, the midpoint of interval [0.684; 1] (see refs) | [0.684; 1] | [4–6] |
| $g$ | Duration of the gonotrophic cycle (in days) | - | $g=50.43-3.202 T+0.054 T^{2}$  Polynomial regression fitted to data in [7,8] | [2.90; 12] | [7,8] |
| $T$ | Local temperature (in Celsius) | - | Field data recorded using data loggers | [15; 32] |  |
| $p$ | Daily survival rate | - | ${p=Par}^{1/g}$ |  | [9] |
| $Par$ | Parity rate: proportion of parous females | - | Field data or predictions of the ‘parity’ GAM model | [0.10; 0.90] |  |
| $n$ | Duration of the extrinsic incubation period (in days) | DENV | $n= 1.03 \times(4 + e^{5.15-0.123 T} )$ |  | [10] |
|  |  | CHIKV | $n= 1244.752 \times\left( e^{-0.215 T} \right)$  regression fitted to data in [11] |  | [11] |
|  |  | ZIKV | $n= 1.03 \times(4 + e^{5.15-0.123 T} )$ |  | [10] |
| $b$ | Infectivity of mosquito to human: probability of vector-to-host transmission | DENV | $b=-3.3705+ 0.2593 T- 0.0043 T^{2}$ | [0; 1] | [12] |
|  |  | CHIKV | 0.49, the midpoint of interval  [0.14; 0.84] (see ref) |  | [13] |
|  |  | ZIKV | $b=-0.81+ 0.0651 T- 0.001 T^{2}$  regression fitted to data in [14] |  | [14] |
| $c$ | Infectivity of human to mosquito: probability of host-to-vector transmission | DENV | 0.31 | [0; 1] | [15] |
|  |  | CHIKV | 0.82, the midpoint of interval  [0.75 ; 0.90] (see ref) |  | [16] |
|  |  | ZIKV | 0.033 |  | [10] |
| $\frac{1}{r}$ | Human infectious period (in days) | DENV | 5, the midpoint of interval  [2 ; 7] (see ref) | [2; 7] | [17] |
|  |  | CHIKV | 4, the midpoint of interval  [2 ; 6] (see ref) |  | [18] |
|  |  | ZIKV | 5 the midpoint of interval  [4 ; 7] (see ref) |  | [19] |

**Supplementary Table S2: Variables selected after bivariate analyses, with justifications for exclusion from multivariate analysis (correlation, VIF, stepwise selection)**

| **Variable topic** | **Variable code** | **Variable name** | **Selected for multivariate analysis (Yes/No)** | **Reason for exclusion** |
| --- | --- | --- | --- | --- |
| Real-time and 48h-lagged microclimatic | rhmax_collection | Maximum relative humidity during sampling | Yes |  |
| Land cover | Patches_low_veget_20 | Number of patches of low vegetation at 20 m | No | VIF>3 |
| Land cover | Edge_High_Veget_100 | Total length of edges of high vegetation at 100 m | No | Correlated with %_High_Veget_20, Environment Residential, Edge_Roads_20, %_Roads_20, Area_Roads_20, %_Buildings_20, Edge_Buildings_20. VIF>3 |
| Land cover | %_High_Veget_20 | Percentage of cover of High vegetation at 20 m | No | Correlated with Edge_High_Veget_100, POP_50, Edge_Roads_20, %_Roads_20, Area_Roads_20, %_Buildings_20, Edge_Buildings_20. VIF>3 |
| Land cover | Area_High_Veget_250 | The average patch size of high vegetation at 250 m | Yes |  |
| Land cover | %_Low_Veget_100 | Percentage of cover of low vegetation at 100 m | No | Correlated with Edge_Low_Veget_100 and Area_Low_Veget_100. VIF>3 |
| Land cover | Edge_Low_Veget_100 | Total length of edges of low vegetation at 100 m | No | Correlated with %_Low_Veget_100 and Environment Residential . VIF>3 |
| Land cover | Area_Low_Veget_100 | The average patch size of low vegetation at 100 m | Yes |  |
| Land cover | Edge_Roads_20 | Total length of edges of roads at 20 m | No | Correlated with Edge_High_Veget_100, %_High_Veget_20, %_Roads_20, Area_Roads_20, Edge_Buildings_20, and Environment Residential . VIF>3 |
| Land cover | %_Roads_20 | Percentage of cover of roads at 20 m | No | Correlated with Edge_High_Veget_100, %_High_Veget_20, Edge_Roads_20, Area_Roads_20, Edge_Buildings_20, and Environment Residential . VIF>3 |
| Land cover | Area_Roads_20 | The average patch size of roads at 20 m | No | Correlated with Edge_High_Veget_100, %_High_Veget_20, Edge_Roads_20, %_Roads_20, Edge_Buildings_20, and Environment Residential . VIF>3 |
| Land cover | Edge_Buildings_20 | Total length of edges buildings at 20 m | Yes |  |
| Land cover | %_Buildings_20 | Percentage of cover of buildings at 20 m | No | Correlated with Edge_High_Veget_100, %_High_Veget_20, Edge_Buildings_20, Area_Buildings_20, Edge_Buildings_20. VIF>3 |
| Land cover | Area_Buildings_20 | The average patch size of buildings at 20 m | No | Correlated with %_Buildings_20. Excluded during stepwise selection. |
| Land cover | Environment (Residential) | Residential environment vs urban parks and built impervious areas | No | Correlated with Edge_High_Veget_100, Edge_Low_Veget_100, Edge_Roads_20, Area_Roads_20, %_Roads_20. VIF>3 |
| Demographic | POP_50 | Population density at 50 m | No | Correlated with %_High_Veget_20. Excluded during stepwise selection. |
| Breeding sites | GL_20 | Number of breeding sites at 20 m | No | Excluded during stepwise selection. |

**Supplementary Table S3: Average estimated R₀ values by environment, with vector abundance and longevity derived from field average (R0_field data)_ or GAM prediction (R0_GAM prediction_), for three arboviruses under three human levels of exposure to bites.** The number of bites/human/day was calculated as the product of the human exposure level and female mosquito abundance in traps per 24h.

| **Environment** | **Virus** | **Human exposure level: 10%** | | | **Human exposure level: 20%** | | | **Human exposure level: 30%** | | |
| --- | --- | --- | --- | --- | --- | --- | --- | --- | --- | --- |
|  |  | **Bites / human / day** | **Mean R0_field data_** | **Mean R0_GAM prediction_** | **Bites / human / day** | **Mean R0_field data_** | **Mean R0_GAM prediction_** | **Bites / human / day** | **Mean R0_field data_** | **Mean R0_GAM predictionn_** |
| Impervious area | CHIKV | 0.45 | 2.03 | 0.54 | 0.90 | 4.07 | 1.08 | 1.35 | 6.10 | 1.61 |
|  | DENV |  | 0.78 | 0.09 | 0.90 | 1.57 | 0.18 | 1.35 | 2.35 | 0.28 |
|  | ZIKV |  | 0.025 | 0.003 | 0.92 | 0.05 | 0.006 | 1.35 | 0.075 | 0.009 |
| Urban parks | CHIKV | 0.81 | 2.43 | 0.80 | 1.62 | 4.86 | 1.60 | 2.42 | 7.29 | 2.40 |
|  | DENV | 0.81 | 0.89 | 0.15 | 1.62 | 1.77 | 0.30 | 2.42 | 2.66 | 0.44 |
|  | ZIKV | 0.81 | 0.03 | 0.005 | 1.62 | 0.06 | 0.01 | 2.42 | 0.09 | 0.015 |
| Residential area | CHIKV | 0.48 | 2.89 | 1.10 | 0.96 | 5.77 | 2.20 | 1.43 | 8.66 | 3.31 |
|  | DENV | 0.48 | 1.15 | 0.26 | 0.96 | 2.31 | 0.52 | 1.43 | 3.47 | 0.79 |
|  | ZIKV | 0.48 | 0.04 | 0.008 | 0.96 | 0.08 | 0.016 | 1.43 | 0.12 | 0.024 |

**Supplementary Table S4: Estimated DENV R₀ values by environment and month derived from field data (R0_field data_) and GAM predictions (R0_GAM prediction_), with 95% confidence intervals, under two human exposure scenarios.** The number of human bites/day was calculated as the product of the human exposure level and female mosquito abundance.

| **Environment** | **Month** | **Human exposure: 0.10** | | | | **Human exposure: 0.20** | | | **Human exposure: 0.30** | | |
| --- | --- | --- | --- | --- | --- | --- | --- | --- | --- | --- | --- |
|  |  | **Number of bites/human/day** | **R0_field data_** | **R0_GAM prediction_** | **95% CI** | **R0_field data_** | **R0_GAM prediction_** | **95% CI** | **R0_field data_** | **R0_GAM prediction_** | **95% CI** |
| Impervious areas | 05 | 0.037 | 0.00 | 0.01 | [0; 0.03] | 0.00 | 0.01 | [0; 0.05] | 0 | 0.01 | [0.00; 0.08] |
|  | 06 | 0.30 | 1.14 | 0.10 | [0.03; 0.41 ] | 2.29 | 0.20 | [0.07; 0.83] | 3.43 | 0.30 | [0.10; 1.24] |
|  | 07 | 1.23 | 0.37 | 0.23 | [0.09; 0.88] | 0.73 | 0.45 | [0.18; 1.74] | 1.10 | 0.68 | [0.27; 2.64] |
|  | 08 | 0.41 | 0.67 | 0.07 | [0.02; 0.27] | 1.35 | 0.13 | [0.05; 0.53] | 2.02 | 0.20 | [0.07; 0.80] |
|  | 09 | 0.58 | 0.87 | 0.14 | [0.05; 0.56] | 1.75 | 0.27 | [0.10; 1.15] | 2.62 | 0.41 | [0.15; 1.67] |
|  | 10 | 0.17 | 2.10 | 0.05 | [0.02; 0.20] | 4.21 | 0.09 | [0.03; 0.40] | 6.30 | 0.14 | [0.05; 0.61] |
|  | 11 | 0.40 | 0.00 | 0.00 | [0.00; 0.01] | 0.00 | 0.00 | [0 ; 0.01] | 0.00 | 0.00 | [0.00; 0.01] |
| Urban parks | 05 | 0.10 | 0.00 | 0.01 | [0.00; 0.03] | 0.00 | 0.01 | [0.00; 0.05] | 0.00 | 0.02 | [0.00; 0.08] |
|  | 06 | 0.34 | 0.65 | 0.13 | [0.05; 0.57] | 1.31 | 0.27 | [0.10; 1.14] | 1.96 | 0.40 | [0.16; 1.72] |
|  | 07 | 2.30 | 2.76 | 0.39 | [0.14; 1.69] | 5.51 | 0.78 | [0.28; 3.37] | 8.26 | 1.17 | [0.42; 5.05] |
|  | 08 | 0.94 | 0.11 | 0.13 | [0.05; 0.52] | 0.21 | 0.25 | [0.10; 1.04] | 0.32 | 0.38 | [0.14; 1.56] |
|  | 09 | 1.17 | 1.20 | 0.23 | [0.09; 1.02] | 2.40 | 0.46 | [0.18; 2.03] | 3.60 | 0.69 | [0.26; 3.05] |
|  | 10 | 0.31 | 1.04 | 0.07 | [0.03; 0.31] | 2.08 | 0.15 | [0.05; 0.63] | 3.13 | 0.22 | [0.08; 0.94] |
|  | 11 | 0.36 | 0.00 | 0.00 | [0.00; 0.00] | 0.00 | 0.00 | [0.00; 0.00] | 0.00 | 0.00 | [0.00; 0.00] |
| Residential areas | 05 | 0.05 | 0.15 | 0.01 | [0.00; 0.08] | 0.29 | 0.03 | [0.00; 0.16] | 0.44 | 0.04 | [0.00; 0.23] |
|  | 06 | 0.34 | 1.08 | 0.27 | [0.10; 1.16] | 2.17 | 0.54 | [0.20; 2.31] | 3.25 | 0.81 | [0.30; 3.47] |
|  | 07 | 1.05 | 2.87 | 0.71 | [0.25; 2.84] | 5.73 | 1.42 | [0.51; 5.68] | 8.60 | 2.13 | [0.76; 8.52] |
|  | 08 | 0.63 | 1.37 | 2.90 | [0.06; 0.79] | 2.74 | 0.37 | [0.12; 1.58] | 4.11 | 0.55 | [0.18; 2.37] |
|  | 09 | 0.67 | 1.86 | 4.37 | [0.14; 1.59] | 3.72 | 0.77 | [0.29; 3.19] | 5.58 | 1.15 | [0.43; 4.78] |
|  | 10 | 0.31 | 0.23 | 1.09 | [0.04; 0.50] | 0.46 | 0.24 | [0.08; 0.99] | 0.69 | 0.36 | [0.13; 1.49] |
|  | 11 | 0.14 | 0.06 | 0.07 | [0.00; 0.03] | 0.12 | 0.02 | [0.01; 0.07] | 0.18 | 0.03 | [0.01; 0.10] |

#### Supplementary figures

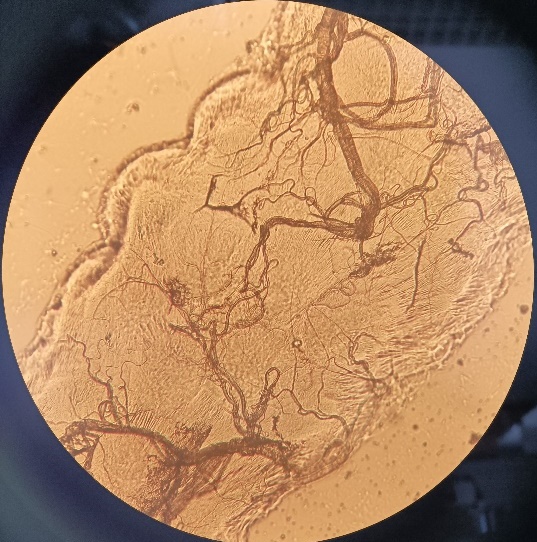

network structure

**A**

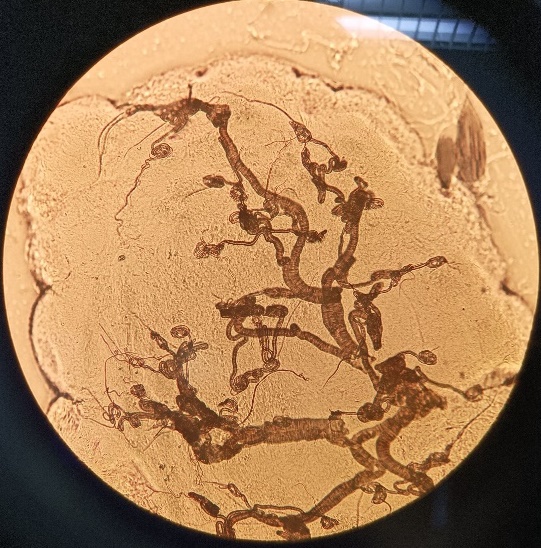

ball formations

**B**

**Supplementary Figure S1: Microscopic image (400× magnification) of tracheoles in the ovaries of (A) parous and (B) nulliparous Ae. albopictus females.** In parous females, the tracheoles form a network-like structure, whereas in nulliparous females, the tracheoles appear coiled in ball-like formations.

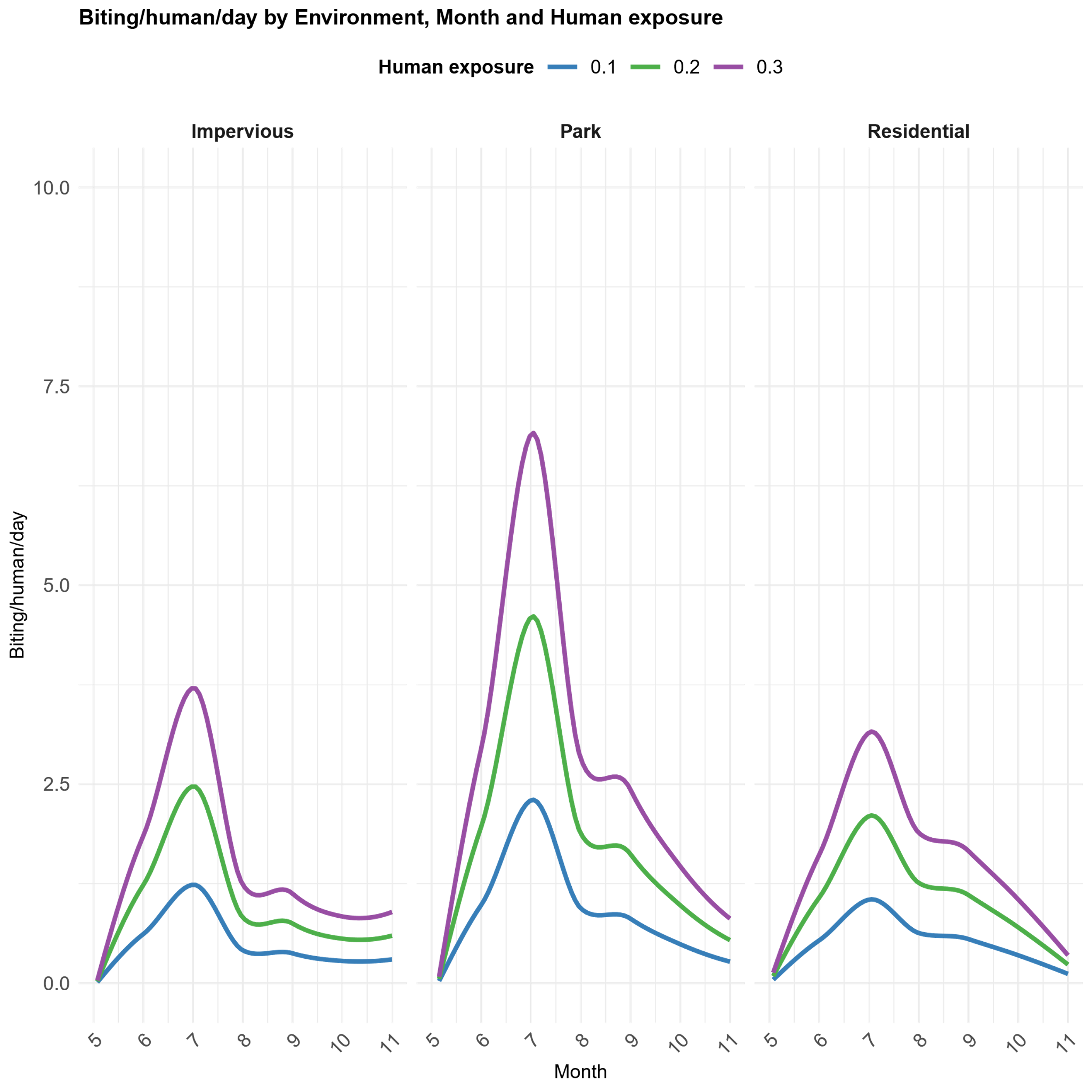

**Supplementary Figure S2: Monthly number of bites per human per day for three different levels of human exposure (10%, 20%, and 30%) across various environments.** The number of bites was calculated by multiplying the number of females per trap per day by the corresponding human exposure level.

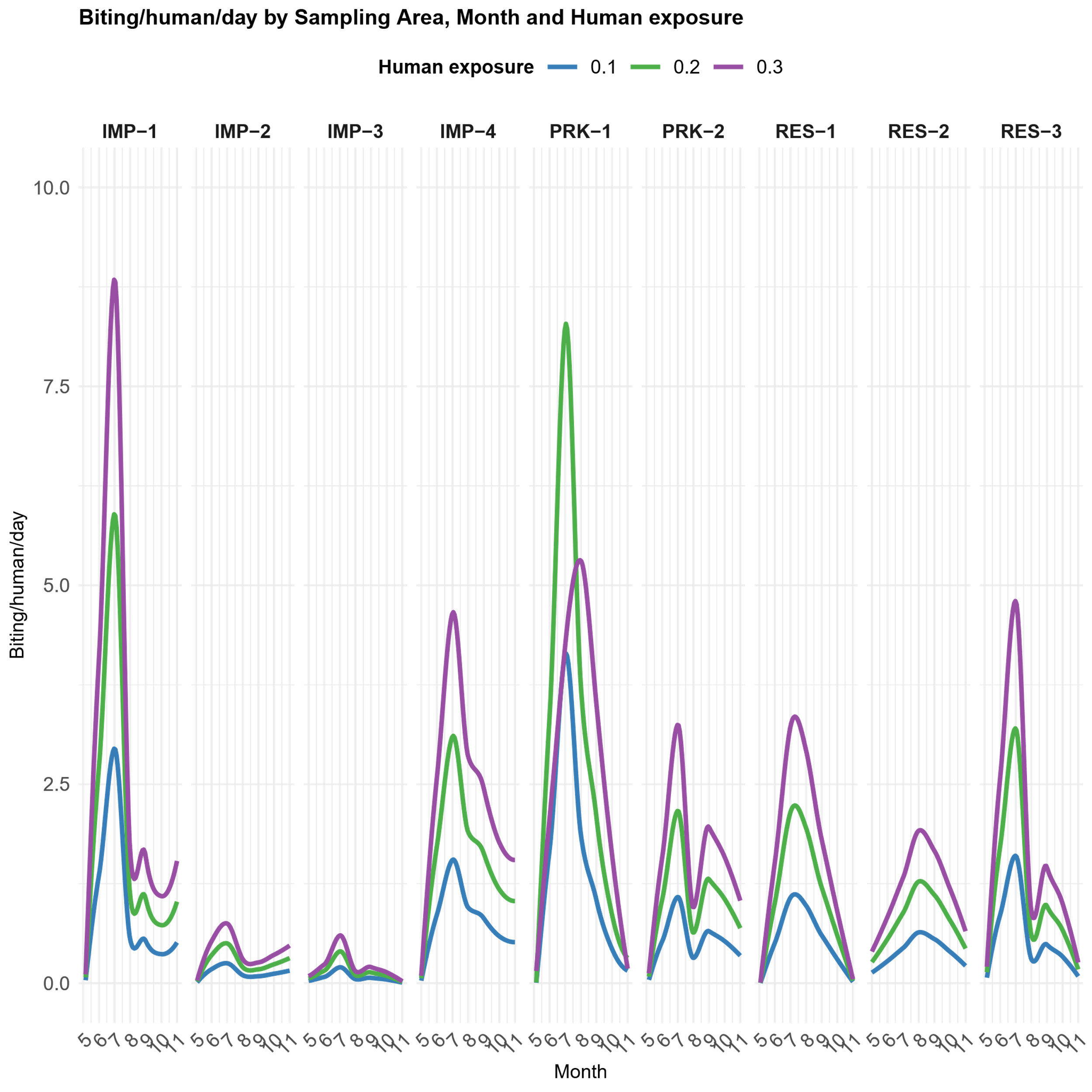

**Supplementary Figure S3: Monthly number of bites per human per day for three different levels of human exposure (10%, 20%, and 30%) across the sampling areas.** The number of bites was calculated by multiplying the number of females per trap per day by the corresponding human exposure level.

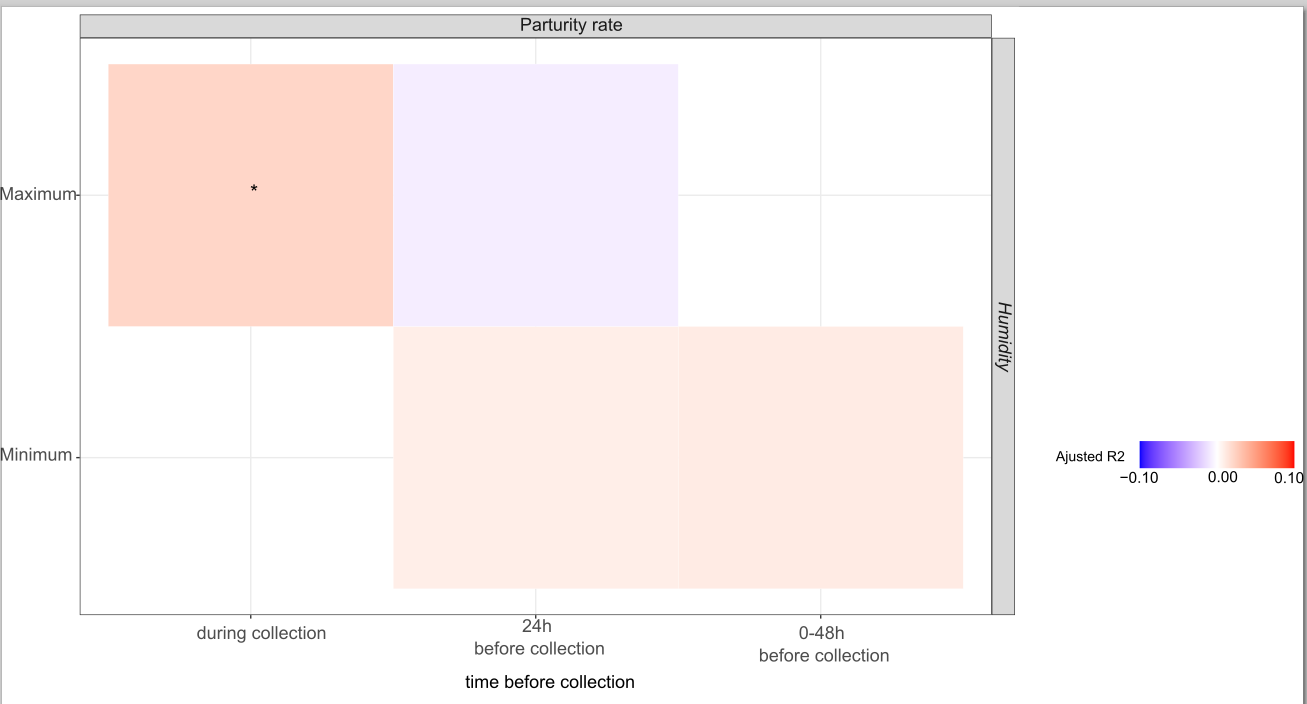

**Supplementary Figure S4: Bivariate relationships between *Ae. albopictus* parity rate and microclimatic variables 48h before and during sampling**. Relationships are analysed based on *Ae. albopictus* parity rate and variables recorded 48h before the start and during sampling sessions. Microclimatic variables include minimum hourly relative humidity (rhmin), maximum hourly relative humidity (rhmax), mean hourly relative humidity (rhmean), minimum hourly temperature (tmin), maximum hourly temperature (tmax) and mean hourly temperature (tmean). The adjusted marginal R² reflects the variance explained by the explanatory variable, adjusted for correlation direction. Boxes are coloured if the p-value is < 0.2 (No asterisk: p-value ∈ [0.05; 0.2], *: p-value ∈ [0.01, 0.05], **: p-value ∈ [0.001; 0.01]; ***: p-value ∈ [0; 0.001]). Box colours depend on the direction of the relationship (blue: negative, red: positive). Colour intensity varies according to the marginal R² value.

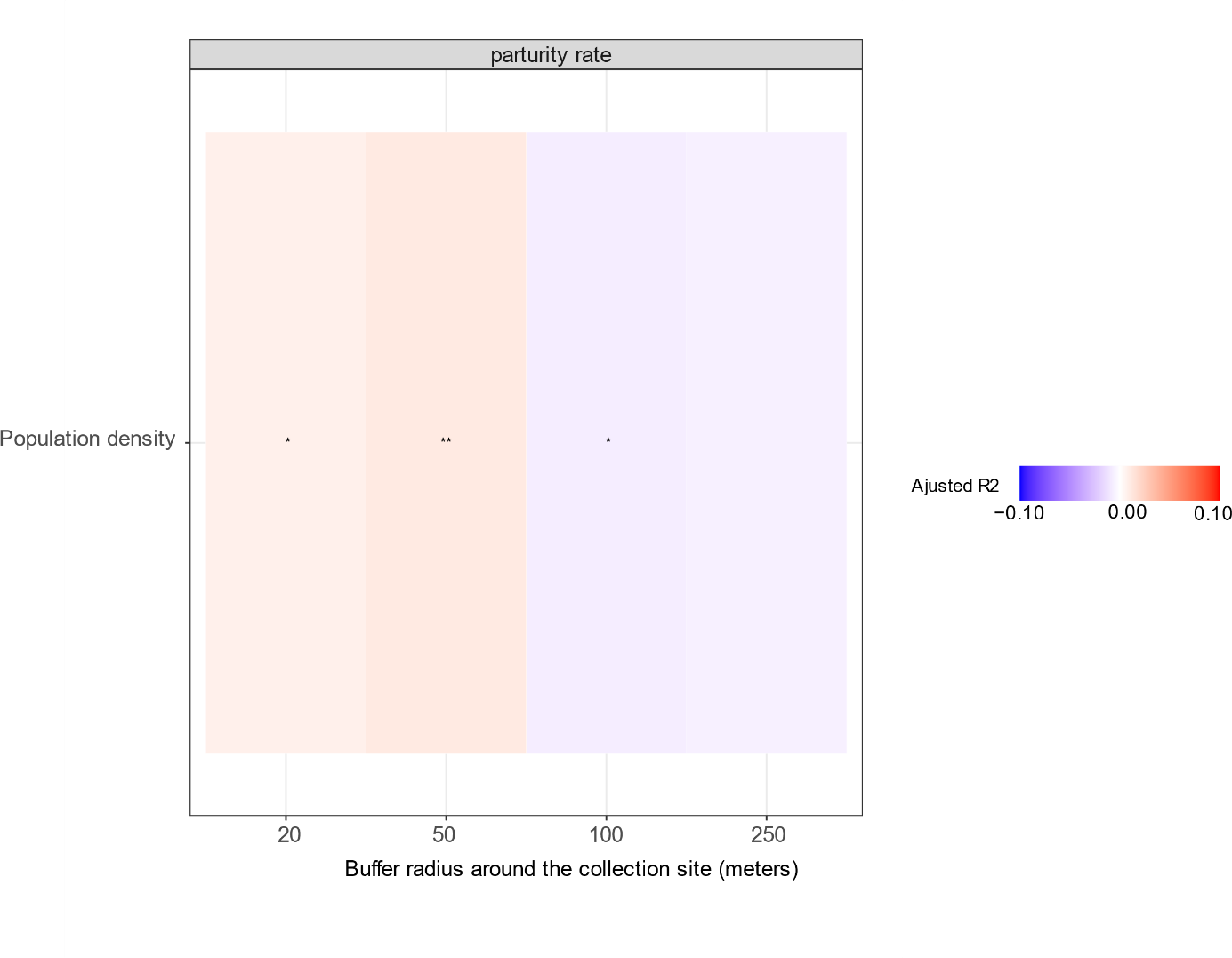
**Supplementary Figure S5: Bivariate relationships between *Ae. albopictus* parity rate and the demographic variables**. The adjusted marginal R² reflects the variance explained by the explanatory variable, adjusted for correlation direction. Boxes are coloured if the p-value is < 0.2 (No asterisk: p-value ∈ [0.05; 0.2], *: p-value ∈ [0.01, 0.05], **: p-value ∈ [0.001; 0.01]; ***: p-value ∈ [0; 0.001]). Box colours depend on the direction of the relationship (blue: negative, red: positive). Colour intensity varies according to the marginal R² value.

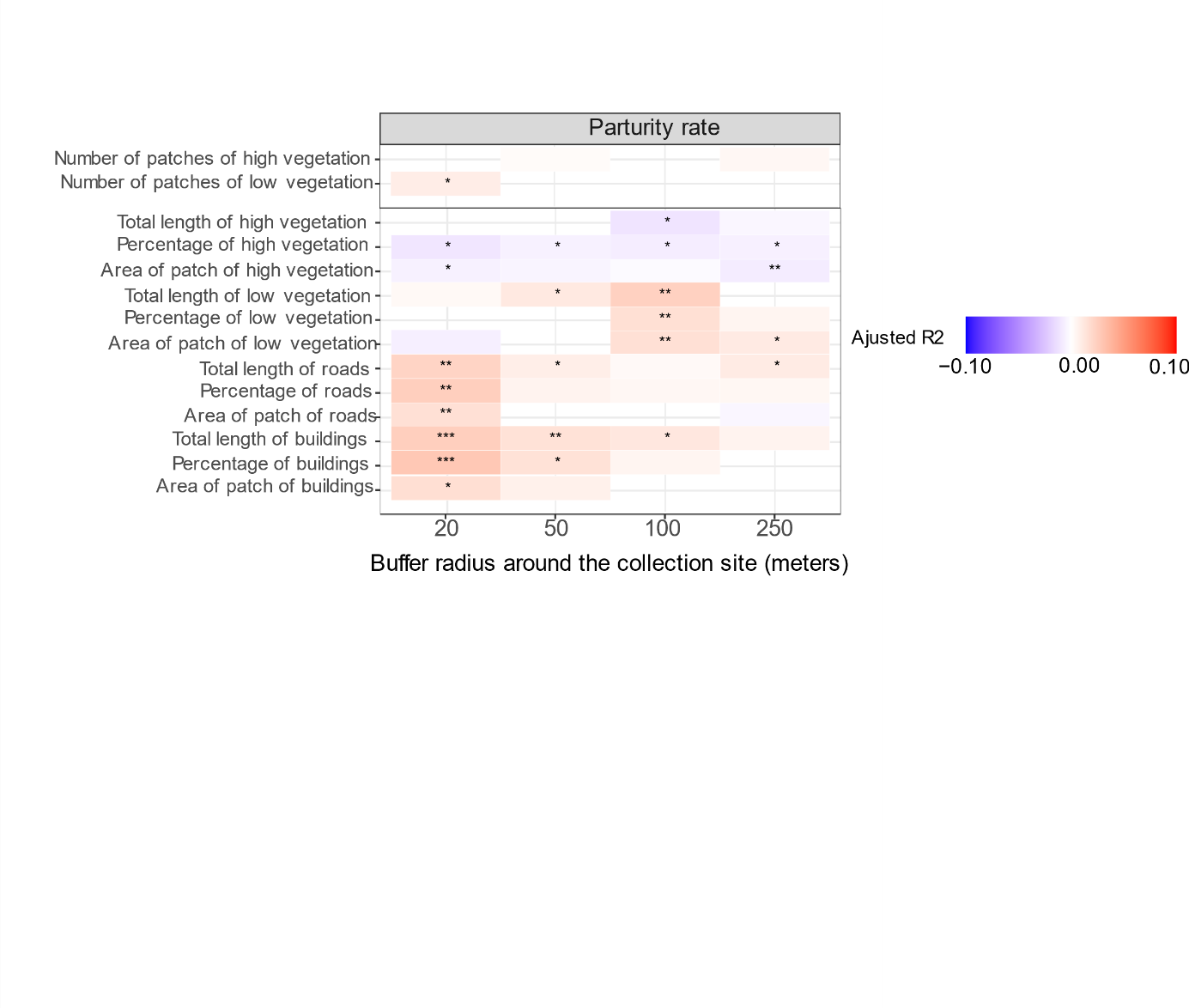

**Supplementary Figure S6: Bivariate relationships between *Ae. albopictus* parity and land cover variables for varied buffer sizes**. Relationships are analysed based on *Ae. albopictus* parity rate and land cover variables for different buffer sizes around traps. The adjusted marginal R² reflects the variance explained by the explanatory variable, adjusted for correlation direction. Boxes are coloured if the p-value is < 0.2 (No asterisk: p-value ∈ [0.05; 0.2], *: p-value ∈ [0.01, 0.05], **: p-value ∈ [0.001; 0.01]; ***: p-value ∈ [0; 0.001]). Box colours depend on the direction of the relationship (blue: negative, red: positive). Colour intensity varies according to the marginal R² value.

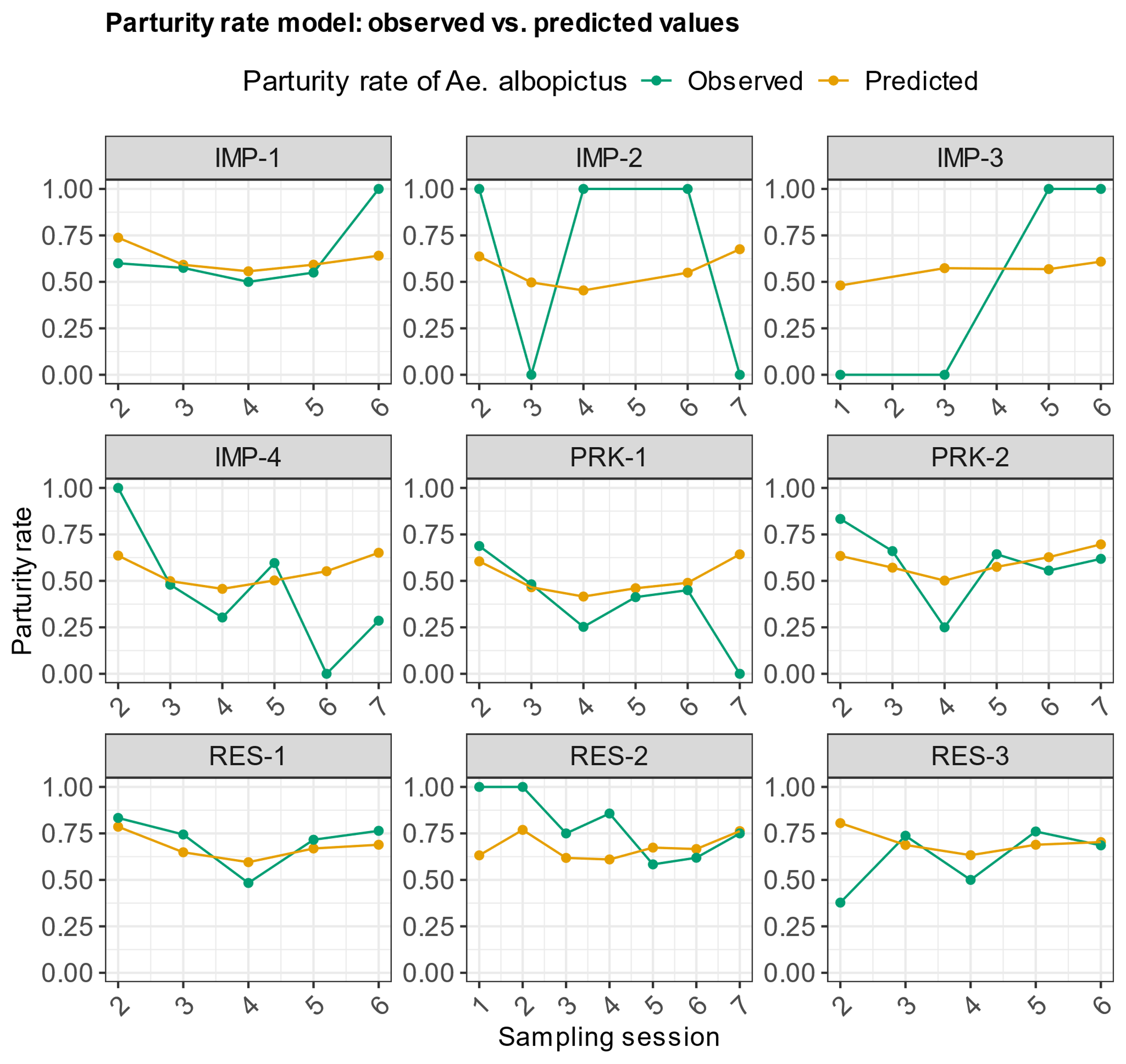

**Supplementary Figure S7: Evaluation of the parity rate model.** The figure compares observed values (green) to predicted values of Aedes albopictus parity rates per area and sampling session. Yellow points represent predictions for each out-of-sample leave-one-area-out cross-validation. The y-axis shows the mean parity rate per area per sampling session, and the x-axis indicates the sampling session.

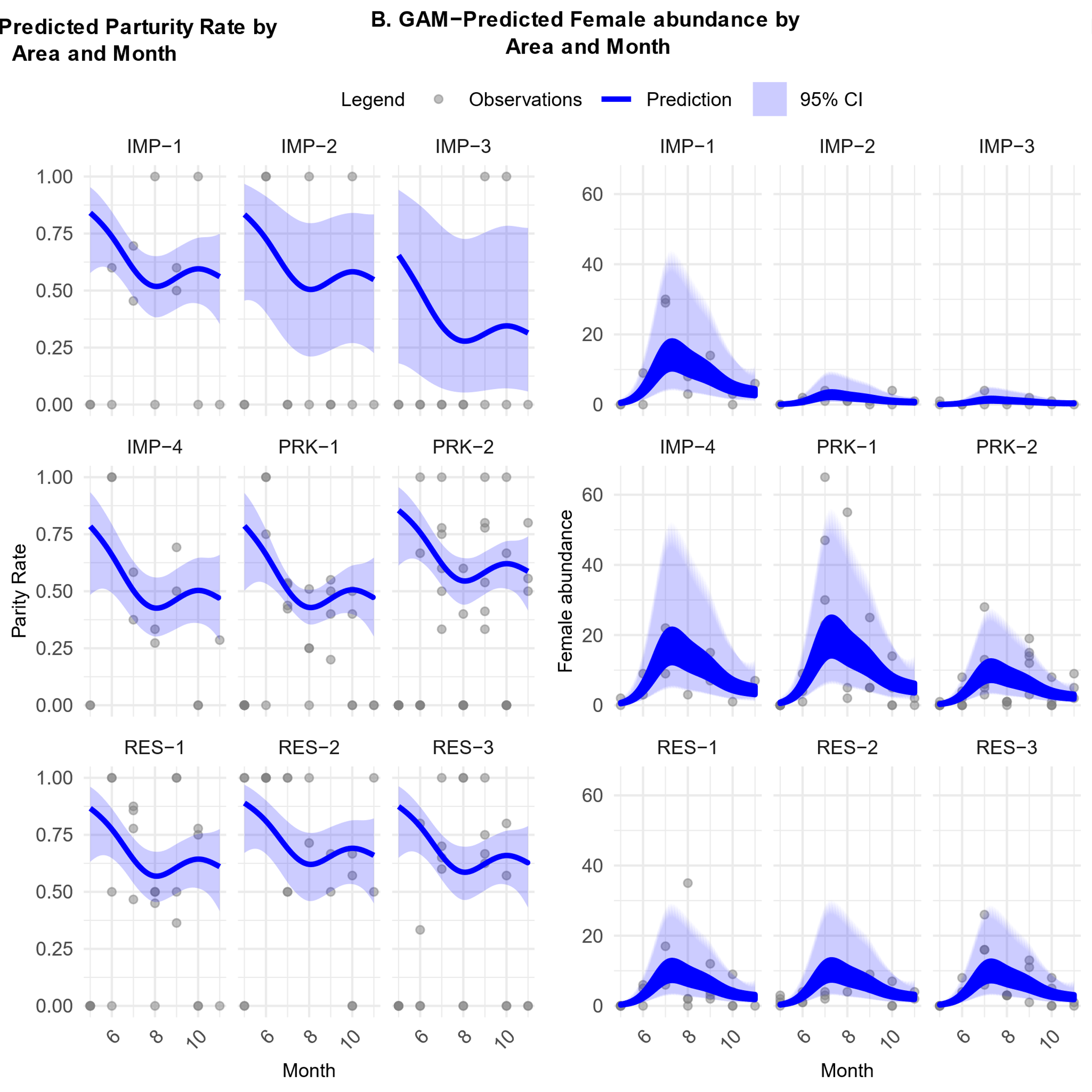

**Supplementary Figure S8: GAM predictions of (A) parity rate and (B) female abundance across sampling areas and months.** The figure shows GAM-based predictions (blue lines) with 95% confidence intervals (blue shading) by area and month, compared to observed values (grey points).

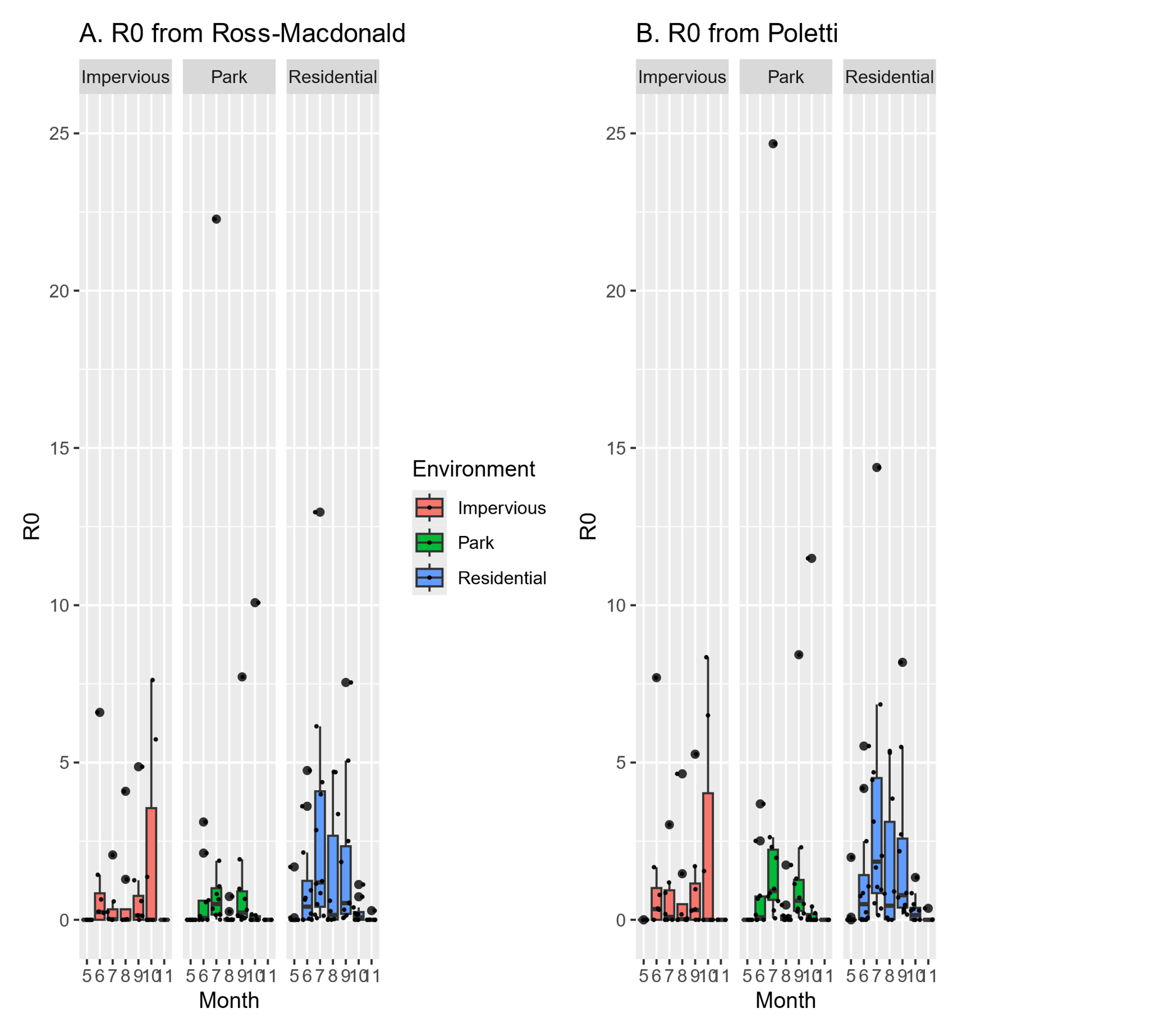

**Supplementary Figure S9: Monthly estimates of DENV R₀ from field data at a human exposure level of 10 %, across different environments.** R₀ was calculated using the Ross-MacDonald formula (A) and the Poletti formula (B).

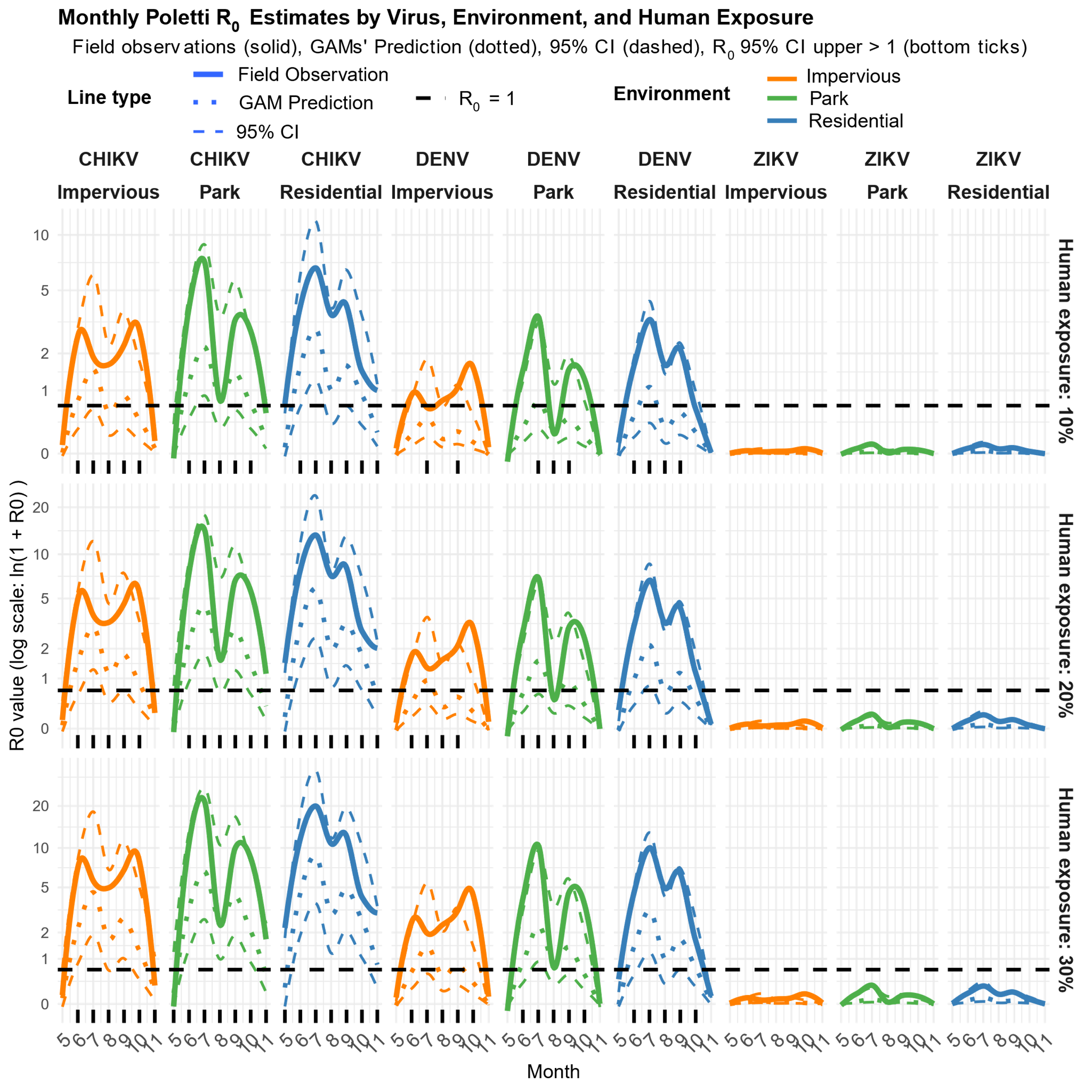

**Supplementary figure S10: Monthly Poletti R₀ estimates based on crude field data and GAM predictions, across different environments, viruses (CHIKV, DENV, and ZIKV), and three levels of human exposure.** The solid line represents R₀ estimates from field data, the dotted line corresponds to GAM-based R₀ predictions, and the dashed lines indicate the 95% confidence intervals. The horizontal black line denotes the epidemic threshold (R₀ = 1). The rug plot along the x-axis highlights months in which the upper limit of the 95% confidence intervals of *R_0_* simulations exceeded 1, and the y-axis is on log-scale.

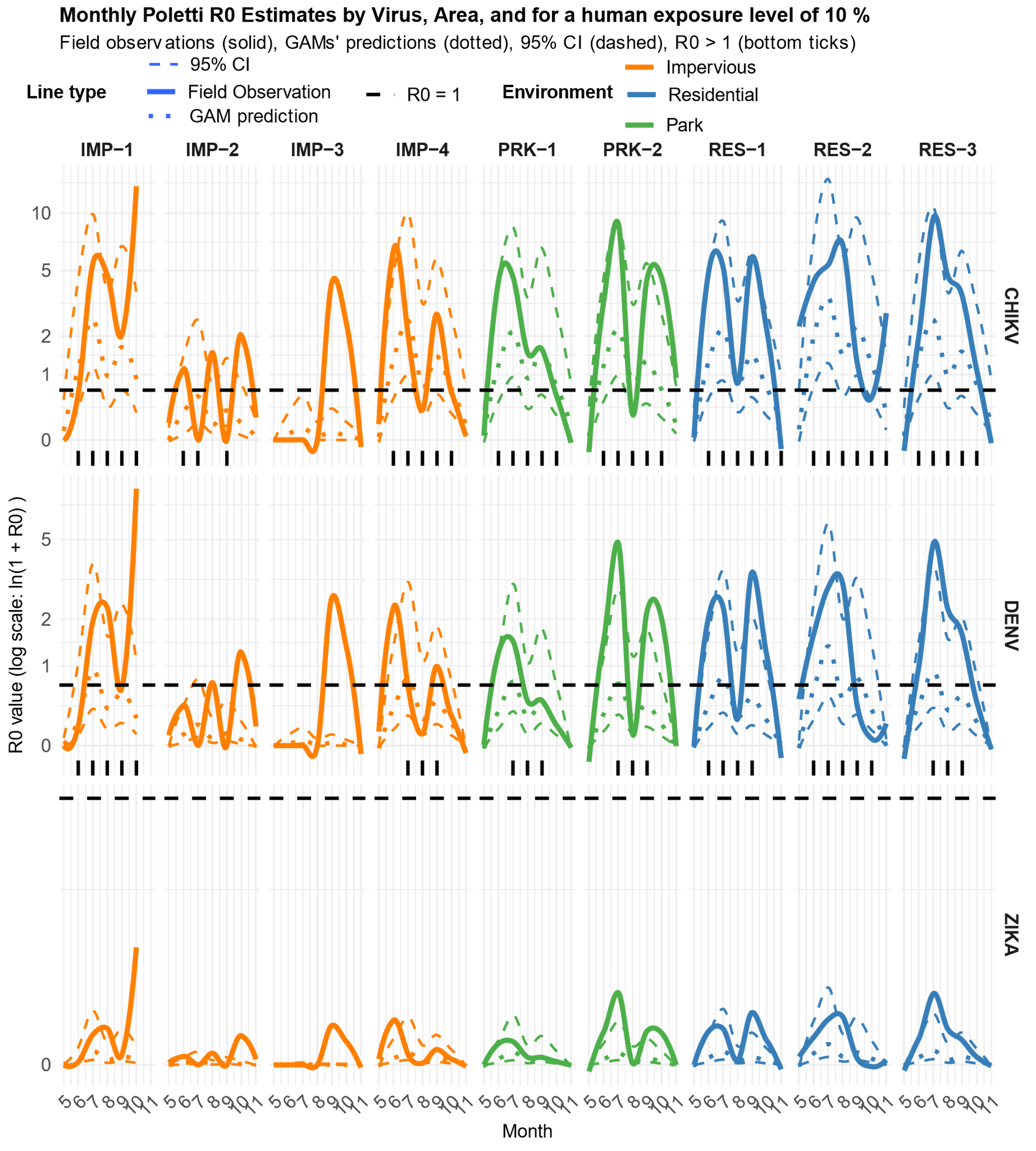

**Supplementary Figure S11: Monthly Poletti R₀ estimates based on field data and GAMs’ simulations across sampling areas and viruses (CHIKV, DENV and ZIKV), with one level of human exposure (10%).** The solid line shows the R₀ estimates from the field data, the dotted line shows the GAM-based R₀ simulations, and the dashed lines show the 95% confidence intervals. The horizontal black line denotes the epidemic threshold (R₀ = 1). The rug plot along the x-axis highlights months in which the R₀ value derived from the field exceeded 1, and the y-axis is on log-scale.

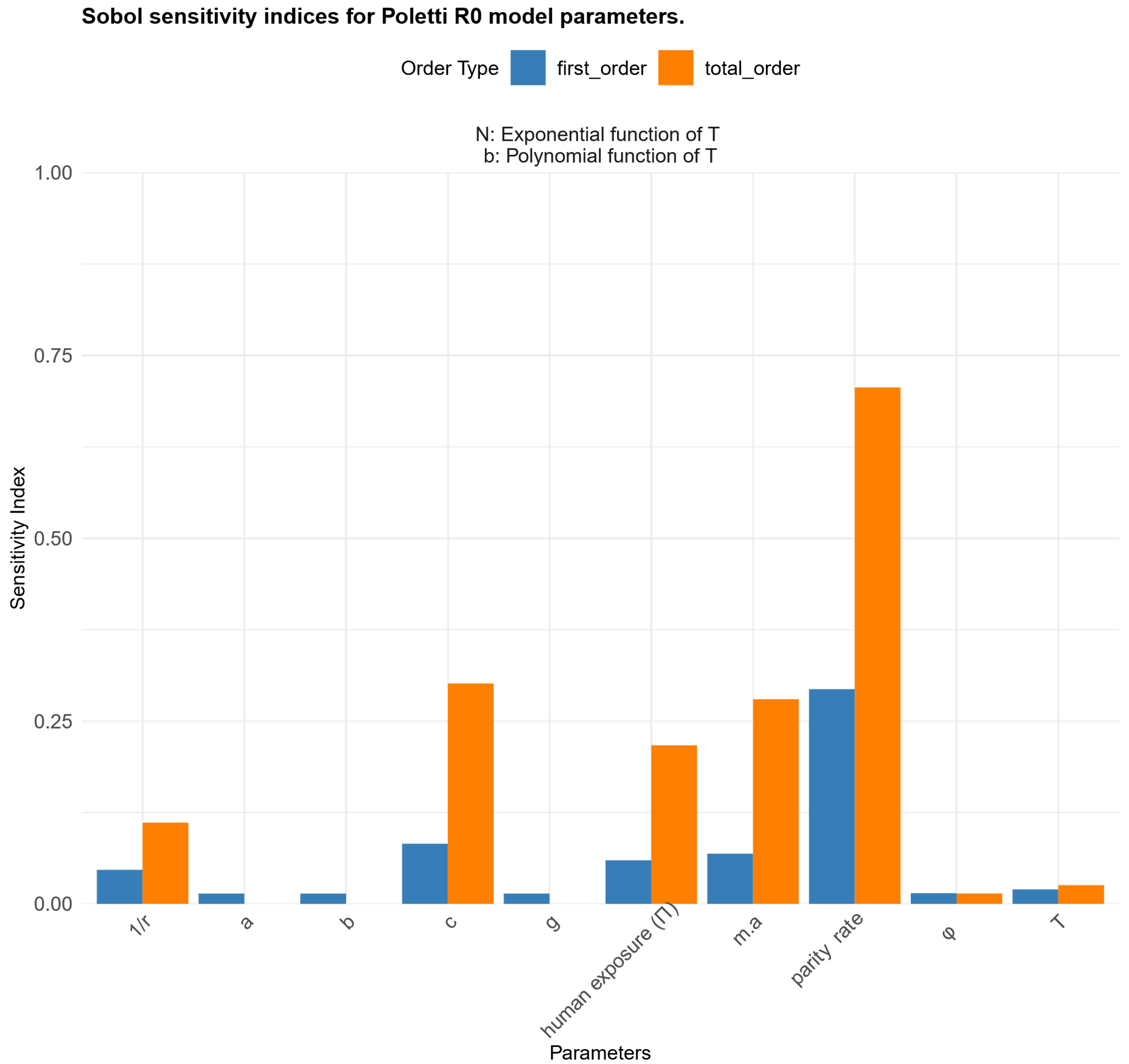

**Supplementary Figure S12: Sobol sensitivity indices for DENV *R_0_* Poletti model parameters, using the formulations of the extrinsic incubation period (n) and vector competence (b) applied in *R_0_* estimation.** Parameters: a = number of blood meals taken from humans per mosquito per day, c = host competence, g = gonotrophic cycle duration, m.a = number of females caught per trap per day, φ= human preference index, T = local temperature, *Π =* human exposure, and 1/r = duration of the human infectious period. First-order and total-order Sobol indices indicate the main and interaction effects of each parameter on the variance of R₀ estimates

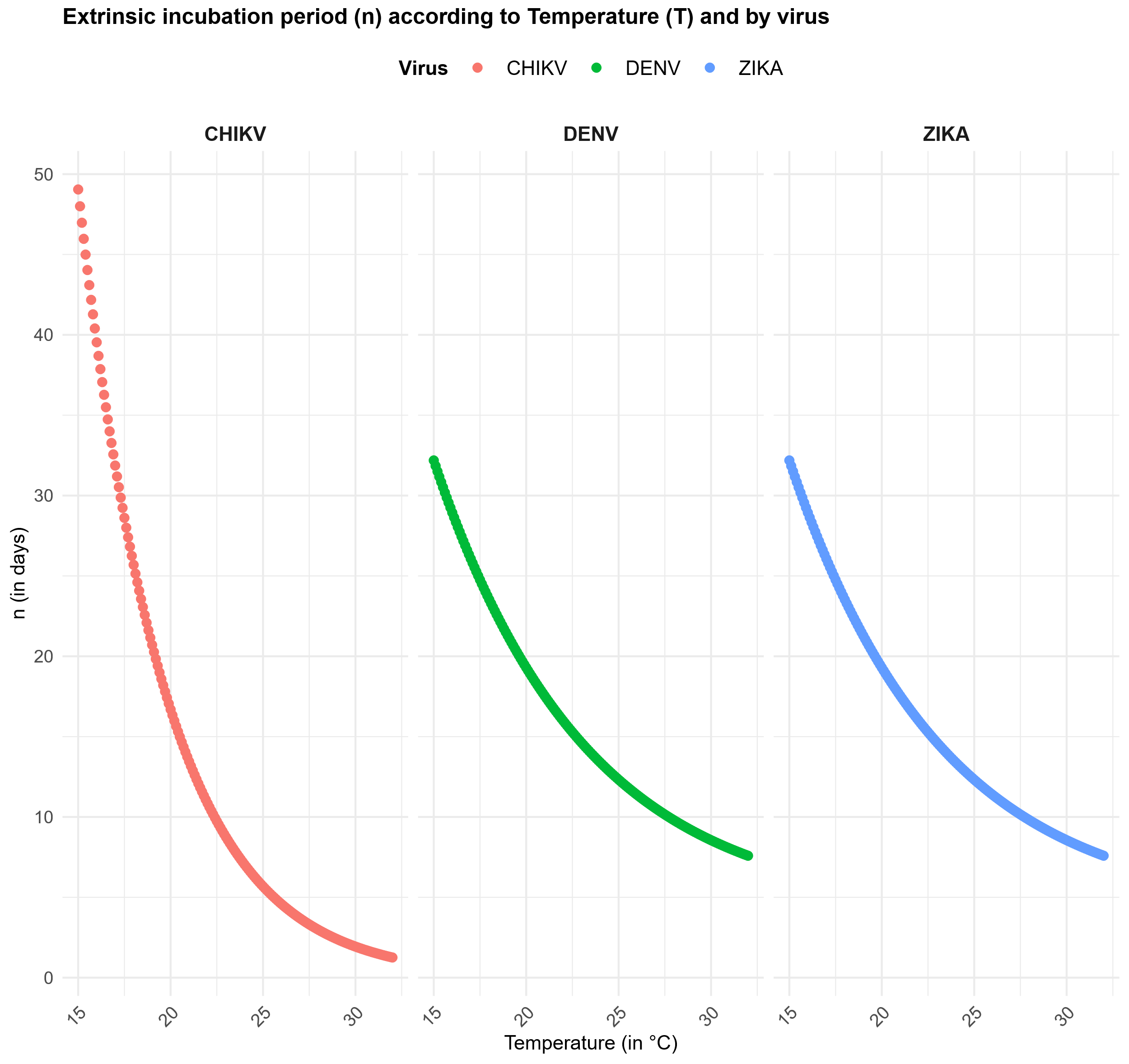

**Supplementary Figure S13: Extrinsic incubation period (n) according to temperature and virus.** Extrinsic incubation period for CHIKV was calculated using the study of Christofferson *et al*. (2023) [10], and for DENV and ZIKV using the formula of Caminade *et al* (2017) [11].

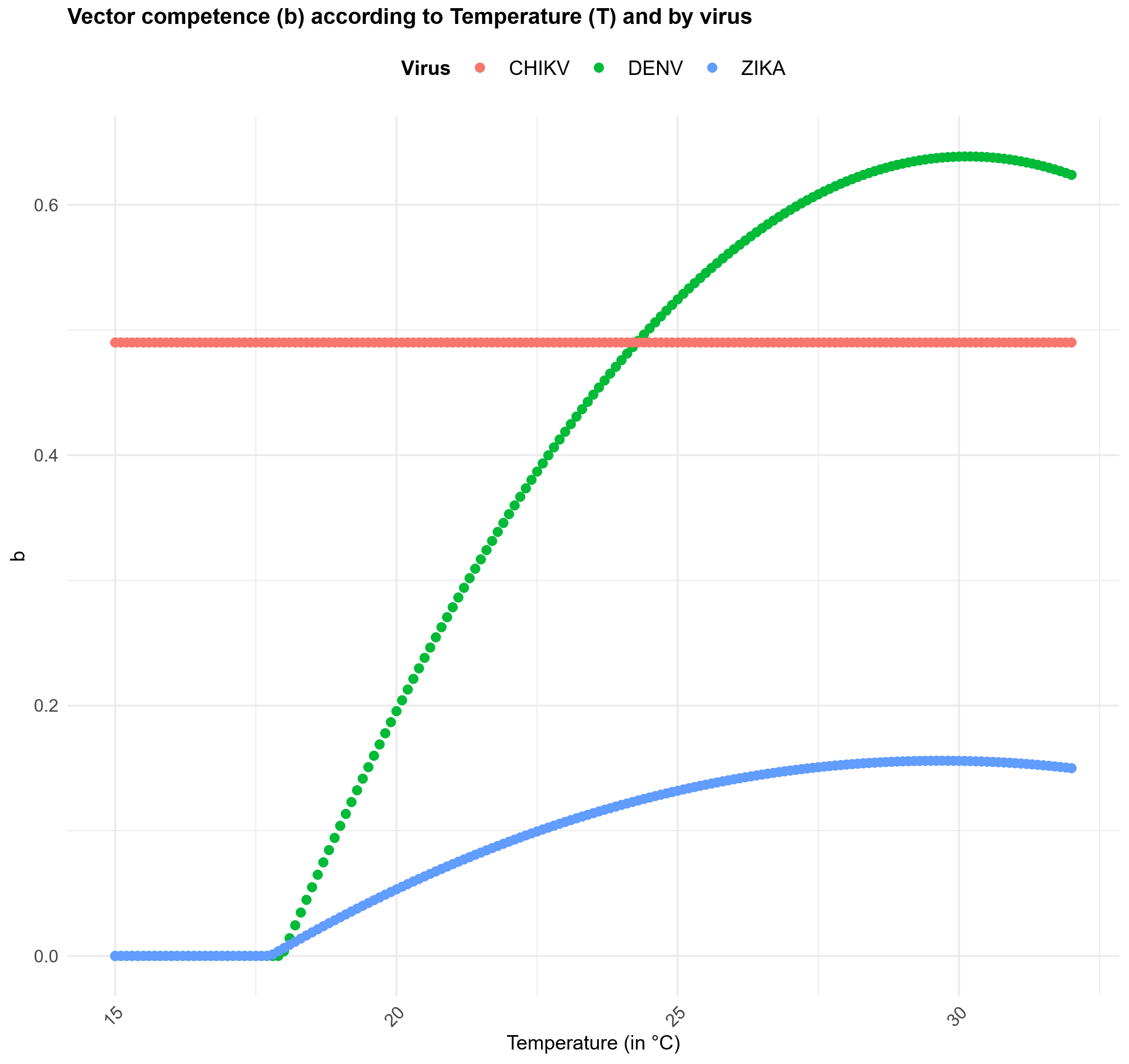

**Supplementary Figure S14: Vector competence (b) according to temperature and virus.** Vector competence for CHIKV was calculated using values of Vega-Rua *et al*. (2013) [13], for DENV using the formula of Benkimoun *et al.* (2021) [14], and ZIKV using the formula of Tesla *et al* . (2018) [15].

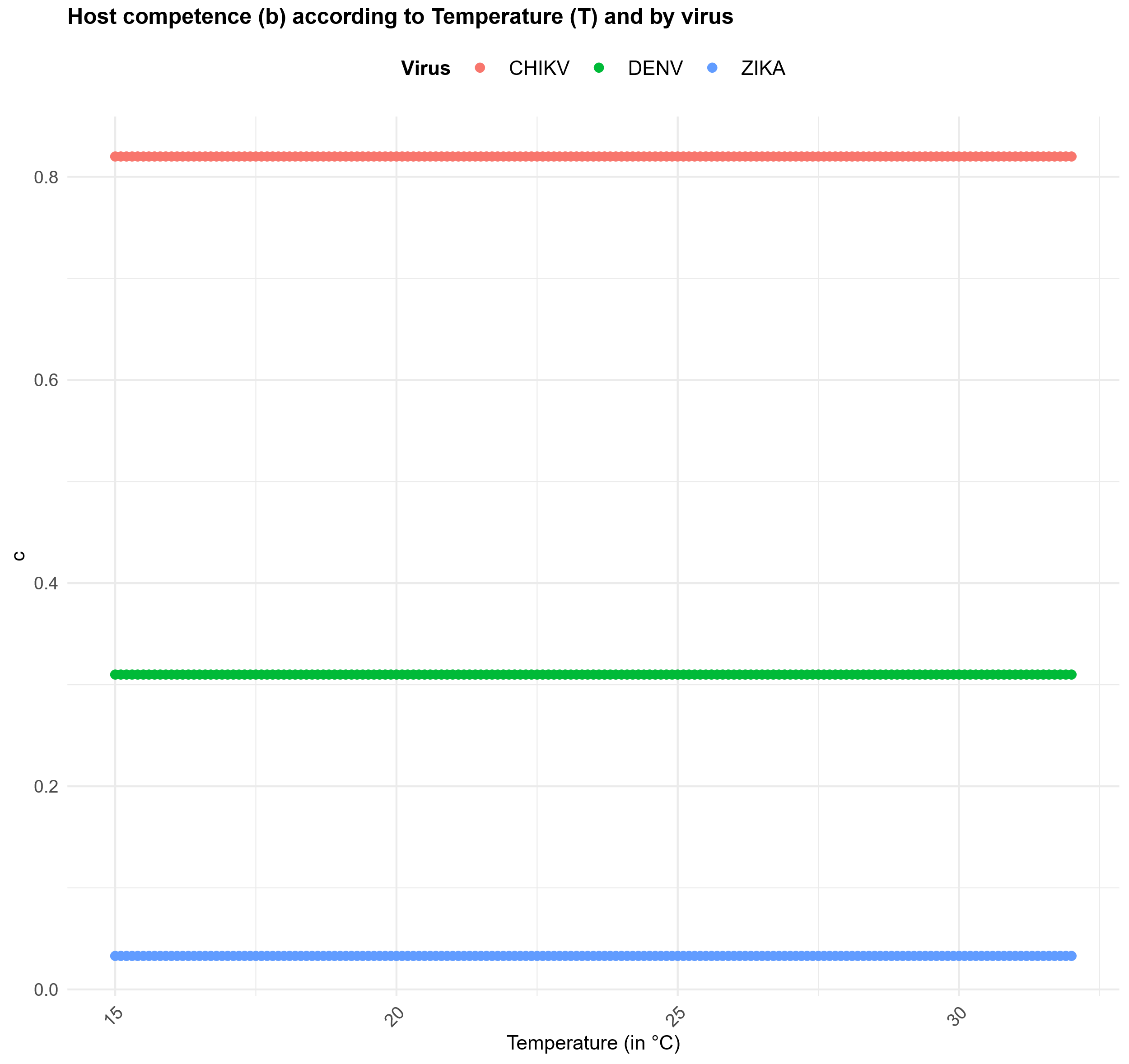

**Supplementary Figure S15: Host competence (c) according to temperature and virus.** Host competence for CHIKV was calculated using values of Metelmann *et al*. (2021) [15], for DENV using the formula of Solimini *et al.* (2018) [16], and ZIKV using the formula of Caminade *et al*. (2017) [10].
